# The potential of Senicapoc, a KCNN4 inhibitor, for the prevention and treatment of breast cancer

**DOI:** 10.1101/2023.04.25.538345

**Authors:** Christos Xiao, Mariska Miranda, Wei Shi, Jonathan Beesley, Jodi M. Saunus, Andrew Civitarese, Debra M. Black, Meagan Ruppert, Melrine Pereira, Susan Jackson, Zachary Teale, Dylan Carter-Cusack, Lauren Kalinowski, Jamie R. Kutasovic, Amy E. McCart Reed, Herlina Y. Handoko, XiaoQing Chen, Darrell Bessette, Kelli MacDonald, Sunil R. Lakhani, Georgia Chenevix-Trench, Kara Britt, Fares Al-Ejeh

**Author notes:** Correspondence Kara Britt, Peter MacCallum Cancer Centre, Melbourne, VIC, 3000, Australia, ORCID 0000-0001-6069-7856, Georgia Chenevix-Trench, QIMR Berghofer, c/o RBH Post Office, Herston, QLD 4029, Australia, ORCID 0000-0002-1878-2587. these authors had equal contribution (Kara Britt, Fares Al-Ejeh). these authors had equal contribution (Christos Xiao, Mariska Miranda, Wei Shi).

## Abstract

**Background:** Genome-wide association studies have identified a breast cancer risk locus at 19q13.31. The candidate causal variants at this locus are located in the first exon of *KCNN4.* KCNN4, which regulates membrane potential and Ca^2+^ signaling, is a good candidate for drug repositioning because its inhibitor, Senicapoc, has been shown to be well tolerated in Phase-II and -III clinical trials for asthma and sickle cell anemia.

**Methods:** We evaluated public mRNA expression data to determine whether the allele at 19q13.31 associated with increased breast cancer risk was associated with *KCNN4* expression. We also used immunohistochemistry to evaluate the relationship between KCNN4 protein expression and breast cancer survival. We then used Senicapoc in two murine mammary tumor models to determine if it would delay tumor development. We also treated mice bearing 4T1 mammary tumors with Senicapoc, by subcutaneous injection and by oral gavage. Finally we used gene editing to make deletions within *Kcnn4* in 4T1 to determine whether Senicapoc had off-target effects on tumor growth.

**Results:** Analysis of the Genotype-Tissue Expression Project showed that the allele at 19q13.31 associated with increased breast cancer risk is associated with increased *KCNN4* expression, suggesting that inhibiting KCNN4 might reduce breast cancer risk. Using immunohistochemistry in a large breast cancer cohort, we found that membrane and cytoplasmic expression is a marker of poor prognosis in triple negative breast cancer. We then tested the efficacy of Senicapoc to prevent and treat breast cancer. This showed that it delays the development of mammary tumors in two murine models, and slows growth of a syngeneic (4T1) model of triple negative breast cancer. Senicapoc monotherapy showed similar efficacy to anthracycline/taxane-based chemotherapy in these studies, with a stronger effect when they were combined.

**Conclusions:** These results provide a rationale for clinical testing of Senicapoc for treating, and even preventing, breast cancer.

## Background

Genome-wide association studies (GWAS) have identified over 200 breast cancer risk loci, including one at 19q13.31 [1]. Fine mapping identified ten candidate causal variants (CCVs) in the first intron of *KCNN4* at the 19q13.31 risk locus. The lead variant (rs56344893) is associated with risk of both estrogen receptor positive (ER^+^) (OR = 1.07 (95% CI 1.05-1.09); P = 2.26E-17) and ER^−^ breast cancer (OR = 1.07 (95% CI 1.05-1.10); P = 6.81E-8) [2]. It is now recognized that drugs which target genes implicated by GWAS are twice as likely to be effective treatments for the studied disease, as exemplified by tamoxifen which targets the estrogen receptor, another GWAS target [3–6]. Several pharmaceutical companies have developed small molecular inhibitors of KCNN4 (recently reviewed in [7]), suggesting the potential for KCNN4 inhibition for the prevention and treatment of breast cancer.

*KCNN4* encodes an intermediate-conductance calcium-activated potassium channel (KCa3.1 or SK4). It is expressed by many cancers and plays an important role in cellular activation, migration and proliferation by regulating membrane potential and Ca^2+^ signaling [8]. Expression of KCNN4 is associated with poor prognosis in breast cancer, particularly in basal-like breast cancer, and higher expression is observed in ER^−^ compared with ER^+^ breast tumors [9], and in chemoresistance in breast tumors [10]. There is also evidence for a pro-tumorigenic role for KCNN4 in hepatocellular carcinoma [11], pancreatic cancer [12], prostate cancer [13], leukemia [14] and glioblastoma [7].

Approximately 15% of breast cancers are triple negative (TNBC), defined as lacking expression of the estrogen receptor, progesterone receptor (PR) and human epidermal growth factor receptor 2 (HER2). Despite their sensitivity to chemotherapy, outcome remains poor for many TNBC patients, particularly in the first five years after diagnosis. Given the limited targeted therapies for this aggressive subtype of breast cancer, there is a need to better understand its biology in order to develop new treatments. Zhang et al [15] showed that suppression of the KCNN4 channel by siRNA and two inhibitors, TRAM-34 and clotrimazole, can inhibit cell proliferation, migration and epithelial-mesenchymal transition in the TNBC cell line, MDA-MB-231, and also promotes apoptosis. There is also evidence that cell cycle control can be modulated by TRAM-34 in murine mammary tumor models [16]. Although it has not yet been tested against TNBC, another KCNN4 inhibitor, Senicapoc (originally known as ICA-17043) is an attractive drug candidate because it has been in Phase III clinical trials, for sickle cell anemia [17] and in two small Phase II clinical trials for asthma [18]. Senicapoc was not found to be efficacious in these trials, but it was well tolerated, is orally bioavailable and has a much longer half-life than TRAM-34 in humans [18].

The purpose of this study was to evaluate the breast cancer risk-associated variants at 19q13.31 and *KCNN4* expression, to assess the relationship between KCNN4 protein expression and outcome in breast cancer, and to test the effect of a KCNN4 inhibitor, Senicapoc, for the prevention of mammary carcinomas in mouse models, and for the treatment of a triple negative murine mammary tumor, 4T1, both as a single agent and in combination with chemotherapy.

## Methods

### Expression quantitative trait loci (eQTL) analyses

GTEx version 8 eQTL summary statistics for "Breast - Mammary Tissue" and "Whole Blood" were retrieved from the GTEx website. Breast cancer GWAS data were downloaded from the Breast Cancer Association Consortium website (https://bcac.ccge.medschl.cam.ac.uk/). Colocalization of expression and risk association signals at 19q13.31 was computed using the HyPrColoc Bioconductor package [19] using default parameters.

### Cell culture

Breast cancer cell lines from ATCC™ (VA, USA), 4T1 and MDA-MB-231, were cultured as per ATCC™ instructions, regularly tested for mycoplasma and authenticated using short tandem repeat profiling. SSM1 and SSM3 mammary carcinoma cells derived from STAT1-deficient mice [20,21] were a kind gift from Robert Schreiber. They were grown in 10% CO_2_ DMEM/F12/10% FBS/1% L-glutamine/1% penicillin-streptomycin/50 μM 2-mercaptoethanol/0.3 μM hydrocortisone/5 μg/mL insulin/10 ng/mL transferrin.

### Quantitative real time polymerase chain reaction (RT-qPCR) for KCNN4

RNA was isolated using the Qiagen RNeasy^®^ kit (Valencia, CA) according to the manufacturer’s instructions. Reverse transcription was performed using Superscript^®^III Reverse Transcription First Strand^TM^ synthesis (Life Technologies, Carlsbad, CA) for human cell lines, and Maxima H Minus First Strand cDNA Synthesis Kit (Life Technologies) for murine cell lines, according to the manufacturer’s instructions. qPCR was performed in an ABI ViiA^TM^ 7 System (Applied Biosystems) with SYBR Green Master Mix (Life Technologies). Data processing was performed using ABI QuantStudio™ Software V1.1 (Applied Biosystems). The quantity of mRNA was calculated using the ΔΔCt method. Three and four primer pairs were designed for human *KCNN4* and murine *Kcnn4* respectively (**Supplementary Table 1**), and the mRNA levels for each sample were measured in technical triplicates for each primer set. Data were plotted using the GraphPad Prism v7.

### Lentiviral knockdown

In order to validate the KCNN4 antibody (clone HPA053841 from Sigma), an MDA-MB-231 breast cancer cell line with *KCNN4* knockdown was generated using a lentiviral approach. Briefly, pLKO.1 plasmids containing shRNAs targeting *KCNN4* and control non-target sequences (**Supplementary Table 1**) were obtained from Sigma-Aldrich (St Louis, USA) and transfected into HEK293T cells with a packaging mix that contains a pCMV-dR8.91 (Delta 8.9) plasmid and a pCMV-VSVG plasmid. Following transfection, fresh medium was added, and cells were incubated for 48 hours for virus production after which the generated virus particles were filtered and used to transduce MDA-MB-231 cells. Following transduction, cell lines with stable expression of *KCNN4* shRNAs were generated by selection in medium containing puromycin at a concentration of 1 μg/mL (Sigma-Aldrich, St Louis, USA).

### Immunofluorescence assays

Cells were grown to approximately 70% confluency on sterilized glass coverslips pre-coated with poly-lysine (Sigma Aldrich^®^) to enhance cell attachment. Coverslips were washed with PBS and fixed in ice cold 4% paraformaldehyde (PFA, Sigma Aldrich^®^) for 15 min at room temperature. Following fixation, cells were washed with PBS and permeabilized in 0.1% Triton X-100 for 10 minutes at room temperature. Coverslips were blocked in filtered 3% bovine serum albumin (BSA, Sigma Aldrich^®^) in PBS overnight. The following day, cells were incubated with primary antibody (15 µg/mL in 3% BSA in PBS): rabbit anti-KCNN4 antibody (IgG, clone HPA053841, Sigma Aldrich^®^) or control rabbit antibody (IgG, 12-370, Millipore) and mouse anti-α-tubulin antibody (IgG, clone DM1A, Sigma Aldrich^®^) for one hour at room temperature. Following primary antibody incubation, cells were washed with PBS and incubated in secondary antibodies (1 µg/mL); Alexa_488_-conjugated anti-rabbit IgG antibody and Alexa_594_-conjuagted anti-mouse IgG antibody (Life Technologies^TM^); and DAPI as a nuclear stain (1:500, D9542 Sigma Aldrich^®^) diluted in blocking buffer for 1 hour at room temperature in the dark. Following secondary antibody incubation, coverslips were washed with PBS and mounted on glass coverslips using ProLong^®^ gold mounting medium (Life Technologies^TM^, Carlsbad, CA). Images of at least four random regions at 60x magnification were acquired using the confocal microscope Zeiss 780 NLO (Carl Zeiss AG, Germany) and were analyzed using Fiji-ImageJ2 [22].

### Immunohistochemistry

KCNN4 protein expression was assessed in the Queensland Follow Up cohort, consisting of 413 formalinLfixed paraffinLembedded breast tumors from patients undergoing surgical resection at the Royal Brisbane Women’s Hospital (RBWH) between 1987 and 1994, with accompanying longLterm (up to 30 years) clinical followLup information [23,24]. The use of clinical samples and patient data for this study was approved by human research ethics committees at The University of Queensland (ref. 2005000785) and the Royal Brisbane and Women’s Hospital (2005/022). Immunohistochemistry (IHC) was performed on 4 μm sections of the tissue microarrays under the following conditions: sections were dewaxed and antigen retrieval performed in citric acid buffer (pH 6.0) at 125 °C for 5 minutes. Exogenous peroxidases were blocked in 3% hydrogen peroxide for 10 minutes followed by blocking in Background Sniper (Biocare Medical) for 15 minutes. Rabbit anti-KCNN4 antibody (clone HPA053841) was diluted 1:50 in Da Vinci Green diluent (Biocare Medical), and the Mach1 kit (Biocare Medical), and incubated with the sections at 4 °C overnight. The Mach1 kit (Biocare Medical) was used for antibody detection. Haematoxylin counterstained sections were scanned at 40x magnification and scored by three observers (J.S. A.M.R. and L.K.). KCNN4 subcellular localization was recorded: cytoplasmic, membrane and nuclear. The maximum score of the duplicate cores were taken for each case and associations between the expression of KCNN4 with clinicopathologic variables and survival were investigated using chi square and log-rank tests (GraphPad Prism v7).

### Mouse studies

All animal work was conducted in accordance with the National Health and Medical Research Council guidelines under the approval of the QIMR Berghofer or Peter MacCallum Cancer Centre Animal Ethics Committees. For dose escalation in non-tumor bearing mice, Senicapoc (20, 40, 80 and 120 mg/kg) was administered in a vehicle of DMSO:oil (1:2) via subcutaneous injection to 8-week old BALB/c mice weekly for 12 weeks. Their body weight was monitored weekly to determine the maximum tolerated dose of Senicapoc.

For the Dimethylbenz[a]anthracene (DMBA) prevention model, Senicapoc (or vehicle control) were administered at 120 mg/kg (3 times a week) via subcutaneous injections for 6 weeks to 8-week old C57BL/6 mice (n = 40 mice/group). The mice were then treated with 1 mg of DMBA via oral gavage weekly for 6 weeks to induce ER^+^ tumors [25] whilst continuing their Senicapoc dosing, and then were monitored for mammary tumors. As expected, some mice developed lymphomas or gastrointestinal tumors prior to the development of mammary tumors and thus were removed from the study.

For the SSM1 and SSM3 prevention models, Senicapoc (or vehicle control) was administered at 120 mg/kg 3 times a week via subcutaneous injections for 6 weeks to 8-week old 129SvEv mice (n = 12/group). The mice were then injected with 100,000 SSM1 or SSM3 cells into the mammary fat pad. The mice were then monitored for mammary tumors and continued treatment with Senicapoc or vehicle.

For treatment models, parental or CRISPR/Cas9-modified 4T1 cells (up to 1 million cells), prepared in PBS:Matrigel (1:1) or RPMI-1640, were injected into the mammary fat pad of 8-week old female BALB/c mice or BALB/c nude mice. Treatments started when the tumors reached up to 75 mm^3^ on average.

For experiments where Senicapoc was given by subcutaneous injection, Senicapoc, Doxorubicin, and Docetaxel were prepared in a vehicle of DMSO:olive oil or corn oil (1:2) for subcutaneous injection on the flanks of the mice. For experiments where Senicapoc was administered by oral gavage, Senicapoc was dissolved in Kolliphor:PEG-400 (1:1) at up to 20 mg/mL and further diluted to the dosage in a final vehicle formulation of Kolliphor:PEG-400:water (1:1:8) [26].

Tumor size and mouse body weight were measured three times a week for assessment of efficacy and toxicity. Tumor growth rates were measured from fitting the tumor growth curves to an exponential growth equation using GraphPad Prism. The percentage change in mouse weights was compared to their body weights on the day when treatment started. The statistical tests were performed using GraphPad Prism.

### Knockout of *Kcnn4* in the 4T1 cell line

4T1 cells were transduced with a lentiviral vector expressing the Cas9 nuclease under blasticidin selection (LentiLCas9L2ALBlast). The Broad Institute’s sgRNA design tool (http://portals.broadinstitute.org/gpp/public/analysis-tools/sgrna-design) was used to design gRNAs. A non-target control sequence (NTC, sgRFP) and three gRNAs targeting mouse *Kcnn4* (*Kcnn4* KO1, *Kcnn4* KO2 and *Kcnn4* KO3, **Supplementary Table 1**) were cloned into a lentiviral vector (lentiGuide-Puro). Cas9-expressing cell lines were transduced with a lentiviral vector expressing NTC or single *Kcnn4* gRNA under puromycin selection.

### Knockout validation by Sanger sequencing and quantitative polymerase chain reaction

After antibiotic selection, cells were harvested and genomic DNA extracted using PureLink Genomic DNA Mini Kit (Invitrogen™). Primers (**Supplementary Table 1**) were designed for PCR amplification of the region flanking the sgRNA targeted regions and for sequencing of the PCR products. Synthego ICE analysis was used to analyze the sequencing results (Synthego Performance Analysis, ICE Analysis. 2019. v3.0. Synthego; [18/05/2021]).

RT-qPCR was performed, as described above, on 4T1 parental control, 4T1 NTC, 4T1 *Kcnn4* KO1, 4T1 *Kcnn4* KO2 and 4T1 *Kcnn4* KO3 to validate the depletion of *Kcnn4* mRNA in knockout (KO) cells. The mRNA levels for each sample were measured in technical duplicates for each *Kcnn4* and housekeeping control primer set (**Supplementary Table 1**).

### Culturing single cell clones

In order to ensure complete *Kcnn4* knockdown, colonies were grown from single cells. 4T1 NTC and 4T1 *Kcnn4* KO3 cell lines were sorted using an AriaIIIu cell sorter, and seeded as single cells into wells of 96-well plates. Once each colony reached confluency in a 6-well plate, genomic DNA was extracted in order to perform sequencing and Synthego ICE analysis. From the Synthego ICE analysis of the single clones (Synthego Performance Analysis, ICE Analysis. 2019. v3.0. Synthego; [16/09/2021]), 12 4T1 NTC and 12 4T1 *Kcnn4* KO3 colonies (all of which had homozygous or compound heterozygous mutations that truncated *Kcnn4*) were selected. Total RNA was isolated from all 24 cell lines as described above. Three primer pairs were used for *Kcnn4*, and two primer pairs were used for *Cas9*, along with two housekeeping genes (**Supplementary Table 1**; CAS9-PR-1 from System Biosciences). The *Kcnn4* and *Cas9* mRNA levels for each sample were measured in technical duplicates for each primer set. For *Cas9* expression, the relative quantitation of each mRNA was normalized to that in the 4T1 parental cells.

### Proliferation of the NTC and *Kcnn4* KO pools

In order to minimize issues relating to heterogeneity in growth rates between individual cells, 12 4T1 NTC and 12 4T1 *Kcnn4* KO3 colonies were pooled to create 4T1 NTC pooled and 4T1 *Kcnn4* KO3 pooled cell lines. 2D and 3D proliferation assays were performed to compare the *in vitro* growth rates of these pooled cell lines. For both assays, five cell lines were compared; 4T1 parental control, 4T1 NTC pre-FACS sort, 4T1 *Kcnn4* KO3 pre-FACS sort, 4T1 NTC pooled cell line and 4T1 *Kcnn4* KO3 pooled cell line. For the 2D proliferation assay, cells were plated in 24-well plates at a concentration of 20,000 cells per well and cultured for 72 hours. For the 3D proliferation assay, a growth in low attachment (GILA) assay was performed and 10,000 cells per well were plated in two 24-well low attachment plates and cultured for 72 hours. At the end of culture a CellTiter Glo assay was performed. For 2D assay cells were carefully washed with PBS three times, and then resuspended in 0.5 mL of CellTiter Glo reagent. For 3D assay 0.1 mL of CellTiter Glo reagent was added to each well and mixed well with the suspended cells. The plate was then placed on a shaker for 10 minutes in the dark. After this, 0.2 mL of Cell Titer Glo/cell suspension was taken and plated onto a 96-well white plate, so that the luminescence signal for each well could be read on a plate reader.

## Results

### Candidate causal variants and KCNN4 expression

We used data from the Genotype-Tissue Expression (GTEx) project, version 8, to determine whether the candidate causal variants at the 19p13.31 risk locus were associated with *KCNN4* expression. None of the variants were associated with expression in normal breast tissue from female donors (*e.g.* rs1685191, P = 0.22, n = 168 donors, Figure 1A), but the risk-associated allele at rs62116961 is associated with increased *KCNN4* expression in whole blood samples from female donors (P = 5.9E-23; n = 254), with colocalizing GWAS and eQTL signals (Posterior Probability = 0.983, Figure 1B). Expression of the main *KCNN4* isoform (ENST00000262888.7) in GTEx breast samples is sex-specific and is significantly higher in younger women (< 50 years of age, P = 0.01, Welch Two Sample t-test; Figure 1C), compared with age- and sex-independent expression in blood (Figure 1D). This observation may have implications for detection of breast cancer-relevant eQTLs in breast tissue, where age-associated changes in gene expression may limit power (< 20% donors in GTEx are under 40 years of age, Figure 1C). Nominally significant associations were observed between the lead variant (rs62116961) and other genes in whole blood (*ZNF155* and *ZNF283*), but there was no evidence of co-localization with the breast cancer risk GWAS signal (Posterior Probability = 0).

**Figure 1.**
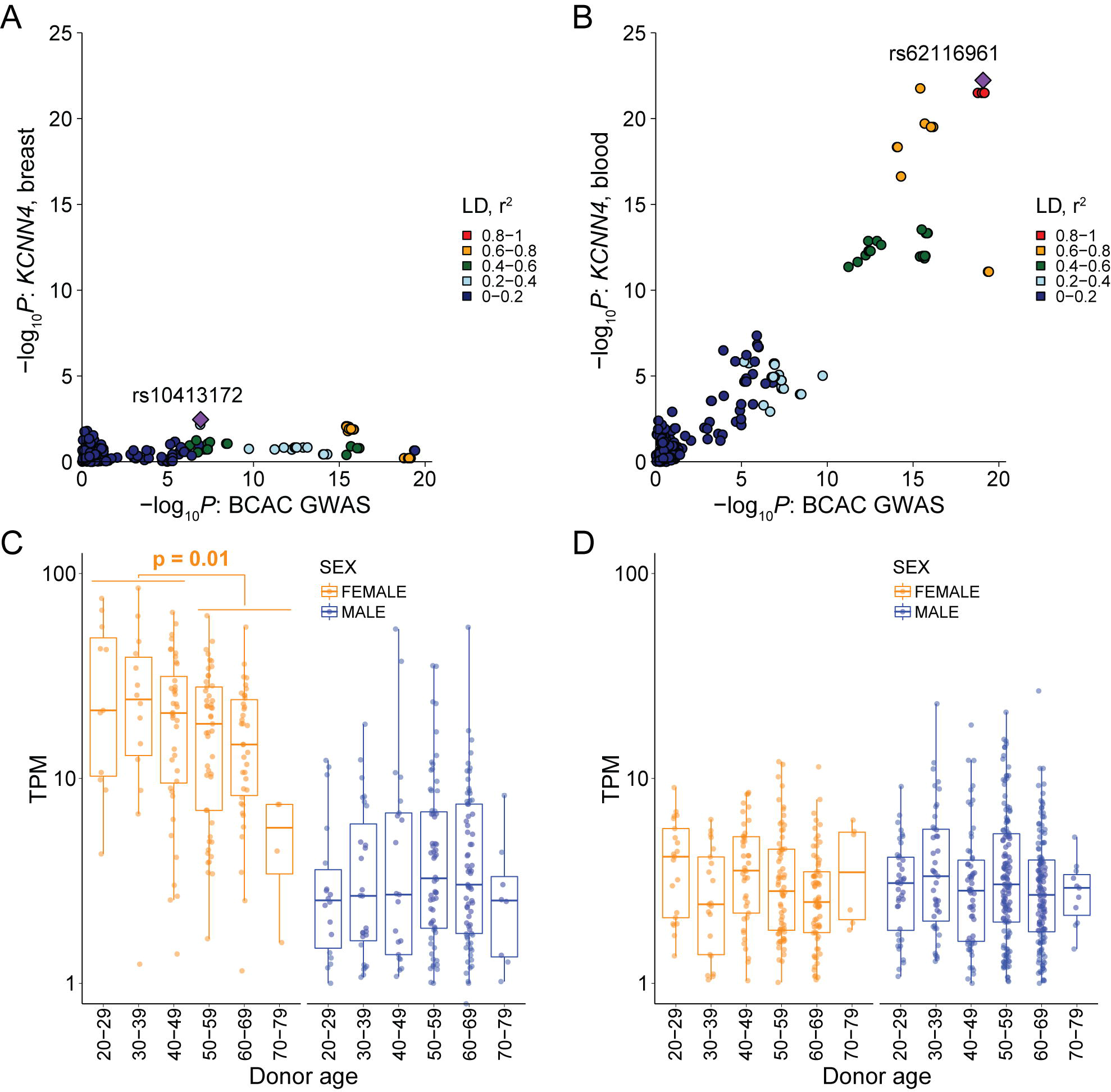
Candidate causal variants and *KCNN4* expression. Association signals for *KCNN4* expression (y-axis) and overall breast cancer risk (x-axis) in breast tissue (**A**); and colocalizing signals in whole blood samples (**B**), using data from female donors in GTEx v8. Colors represent pairwise LD r^2^ coefficients with the top eSNP (labeled purple diamond), based on 1000 Genomes Project Phase 3 version 5 data. Expression levels of the canonical *KCNN4* transcript stratified by age and sex in GTEx breast (**C**) and whole blood (**D**) samples. Individual samples are represented by dots and box plots show median expression (50th percentile; middle line), interquartile range (IQR; 25th to 75th percentiles; box edges) and distribution tails (whiskers).

### KCNN4 mRNA expression in breast cancer

Next, we interrogated *KCNN4* expression in The Cancer Genome Atlas (TCGA) human breast cancer RNA-seq data. *KCNN4* mRNA levels were significantly higher in TNBC compared to other immunohistochemistry (IHC) subtypes, and in basal-like breast cancer (BLBC) compared to other intrinsic (Prediction Analysis of Microarray, PAM50) subtypes (Figure 2). While *KCNN4* expression was not associated with survival in all cases of breast cancer in the TCGA dataset (Figure 3), high *KCNN4* expression was associated with poorer survival in patients across all subtypes treated with chemotherapy, particularly in patients treated with chemotherapy without radiotherapy (Figure 3). There was no relationship between *KCNN4* expression and survival of patients treated with endocrine therapy, with or without chemotherapy, nor of patients treated with chemotherapy and radiotherapy (**Supplementary Figure 1**).

**Figure 2.**
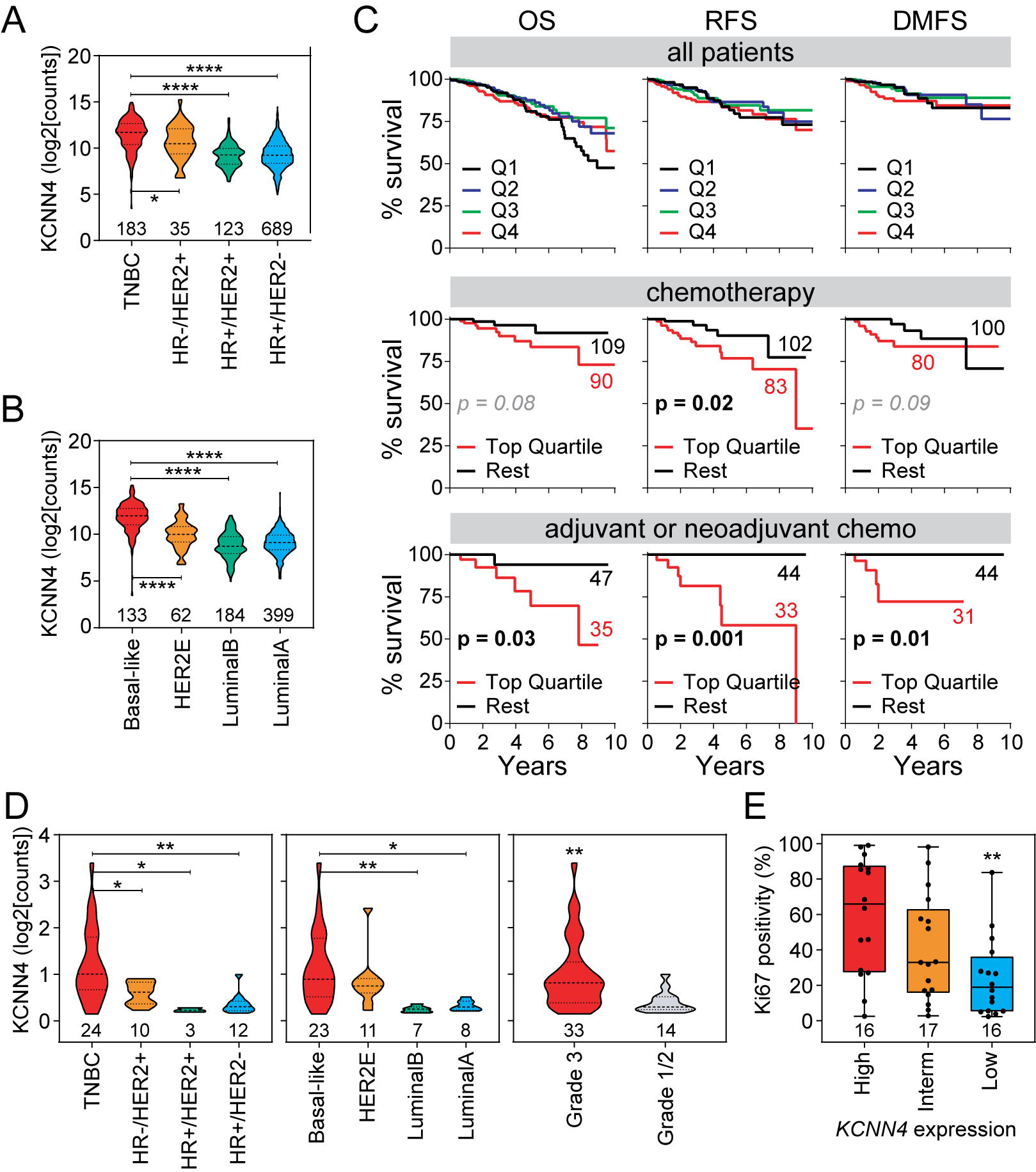
*KCNN4* expression in breast cancer. Expression of *KCNN4* mRNA in the TCGA breast cancer RNA-Seq dataset, comparing IHC subtypes (**A**) or intrinsic (PAM50) subtypes (**B**). One-way Anova P values shown (* P < 0.05, *** P < 0.001, **** P < 0.0001). Case numbers are shown for each of the groups.

**Figure 3.**
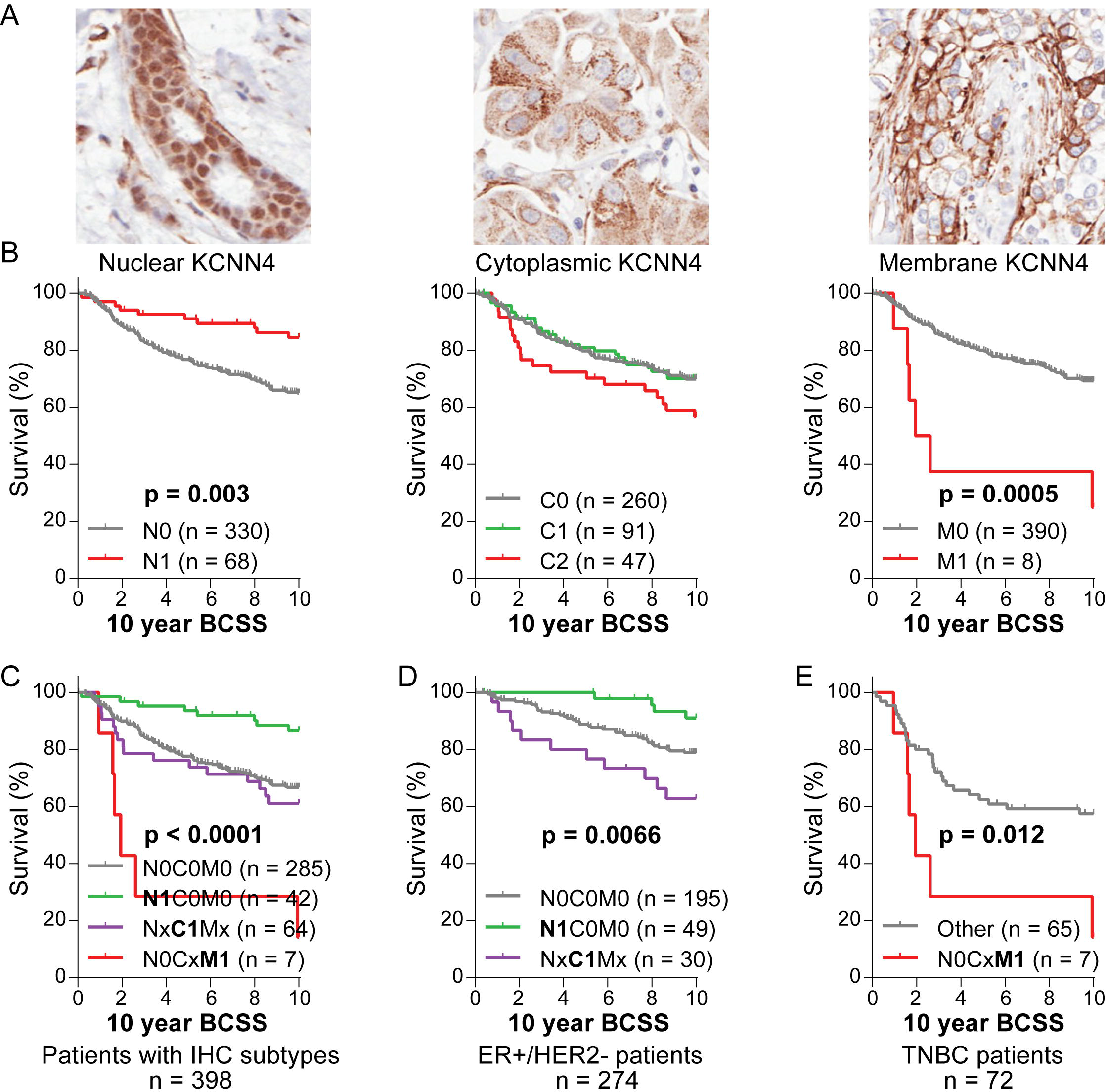
High *KCNN4* associates with shorter survival after chemotherapy. Overall survival (OS), relapse-free survival (RFS) or distant metastasis-free survival (DMFS) of TCGA breast cancer cases were compared after categorizing cases by treatment regimen (chemotherapy but no radiotherapy) and *KCNN4* expression quartiles (Q1: bottom; Q2/Q3: middle quartiles; Q4: top quartile). Log-rank test was used for statistical analysis and significant or near significant P-values are reported.

Because these findings implicate *KCNN4* in chemotherapy resistance, we investigated its expression in residual breast tumor tissue from women who had incomplete responses to neoadjuvant chemotherapy (NAC) [27]. *KCNN4* expression in residual tumors after chemotherapy was significantly higher in TNBC and BLBC cases compared to all other subtypes, and in residual tumors of high grade (Figure 4A). Proliferative activity in post-NAC residual tumor cells is a marker of long-term outcome [28], and we found a direct relationship between *KCNN4* and the proliferation marker Ki67 in these samples (Figure 4B), further implicating KCNN4 in aggressive clinical behavior.

**Figure 4.**
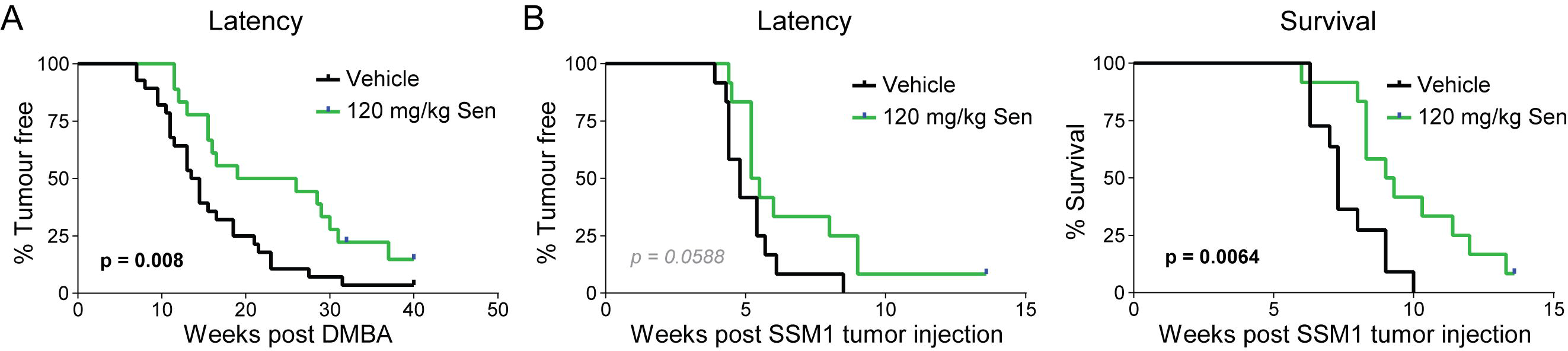
Expression of *KCNN4* mRNA in post-neoadjuvant chemotherapy residual breast tumors. (**A**) *KCNN4* mRNA digital counts were compared across IHC subtypes (left panel), intrinsic (PAM50) subtypes (middle) using One-way Anova (* P < 0.05, ** P < 0.01), and high versus low-intermediate grade tumors (right) using student’s t-test (** P < 0.01). (**B**) These residual breast tumors were classified according to *KCNN4* mRNA expression terciles and compared for Ki67 positivity (one-way Anova was used for statistical comparisons ** P < 0.01). Case numbers are shown for each of the groups. Data originally from [27].

### KCNN4 protein expression in breast cancer

To assess KCNN4 protein expression and survival in breast cancer patients, we selected a rabbit antibody against human KCNN4 (clone HPA053841 from Sigma) as this affinity purified antibody is specified for IHC use and was used in the Human Protein Atlas [29] (http://www.proteinatlas.org). We evaluated *KCNN4* expression in human breast cancer lines by RT-qPCR (**Supplementary Figure 2**) and selected MDA-MB-231 for antibody validation. Using MDA-MB-231 cells expressing shRNAs against *KCNN4*, we confirmed the specificity of the KCNN4 antibody by immunofluorescence and IHC (**Supplementary Figure 3A and B**).

We then performed IHC analysis of KCNN4 expression in the Queensland follow-up cohort [23,24], a consecutive series of 413 breast cancer cases of which 398 had associated ER/PR/HER2 IHC subtype annotations. We observed frequent KCNN4 positivity in endothelial cells (**Supplementary Figure 3C**) of tumor vasculature (272 (66%) cases with such cells present) but this did not associate with any of the clinicopathological indicators (**Supplementary Table 2**). Within the cancer cells, irrespective of vasculature staining, KCNN4 staining in tumor cells (Figure 5A) was observed in 190/413 cases (46%). Nuclear staining in tumor cells was observed in 73/413 cases (17.7%); 143/413 cases (34.6%) had granular cytoplasmic staining; either as strong (50 cases, 12.1%) or weak (93 cases, 22.5%) and 8/413 cases (1.9%) had focal membrane staining. KCNN4 membrane staining in tumor cells was significantly associated with higher tumor grade, ER negativity and the TNBC subtype (**Supplementary Table 2**). Strong cytoplasmic KCNN4 staining in tumor cells was significantly associated with high tumor grade, whereas nuclear staining was significantly associated with ER positivity and higher proliferation (**Supplementary Table 2**). Nuclear KCNN4 staining was significantly associated with better overall survival (P = 0.003), and membrane KCNN4 staining was significantly associated with poor survival (P = 0.0005; Figure 5B).

**Figure 5.**
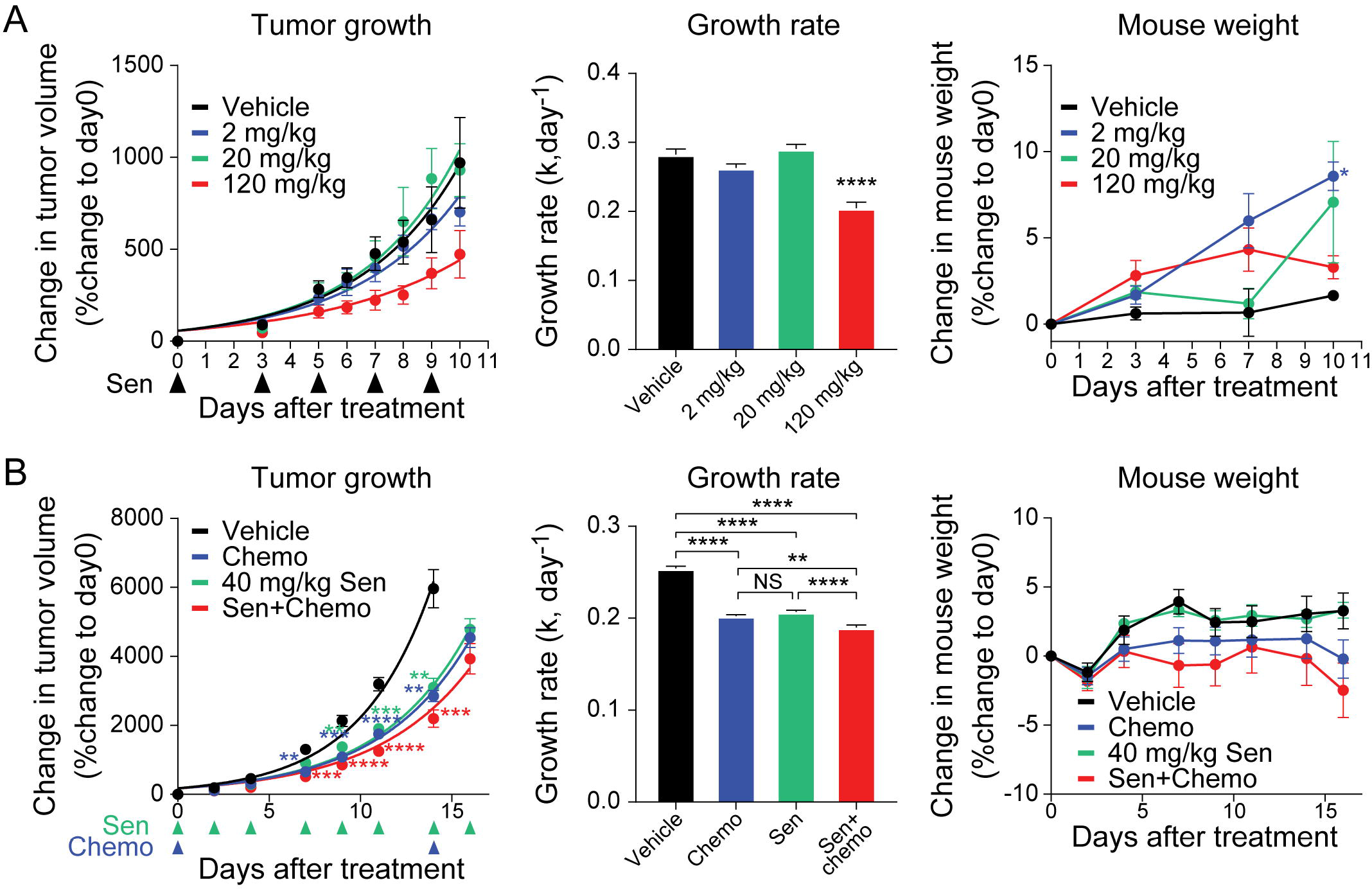
Subcellular compartment-specific expression of KCNN4 in human breast tumors. (**A**) Representative IHC images showing patterns of KCNN4 staining in tumor cells: nuclear, strong cytoplasmic and membrane. Subcellular staining was scored as: nuclear negative (N0) or positive (N1), cytoplasmic negative (C0), weak (C1) or strong (C2), and membrane negative (M0) or positive (M1). TMA cores are 0.6 mm^2^ and magnified insets are 125 μm^2^. (**B**) Kaplan Meier analysis of the association between KCNN4 subcellular expression and 10-year breast cancer-specific survival (BCSS). Log-rank P values shown. (**C-E**) KCNN4 expression patterns were consolidated into four categories in analyses stratified by IHC subtypes; N0C0M0: negative in all cell compartments, **N1**C0M0: exclusive nuclear staining, Nx**C1**Mx: strong cytoplasmic staining irrespective of nuclear or membrane staining, and N0Cx**M1**: membrane positive staining in the absence of nuclear staining. Total case numbers and case numbers in each group in panels **B**-**E** are shown, and statistical significance was determined with log-rank test.

Given the association of KCNN4 staining in the tumor cells with clinicopathological indicators and survival, we consolidated the KCNN4 staining as membrane positive (M1) or negative (M0), strong cytoplasmic positive (C1) or weak/negative (C0), nuclear positive (N1) or negative (N0). We then further consolidated the KCNN4 staining patterns in the different subcellular localization into four groups: N0C0M0 was negative for all cell compartments, **N1**C0M0 cases had nuclear staining in the absence of strong cytoplasmic and membrane staining, Nx**C1**Mx had strong cytoplasmic staining irrespective of nuclear or membrane staining, and N0Cx**M1** consisted of cases with membrane positivity without nuclear positivity and irrespective of cytoplasmic staining. The consolidated staining patterns associated significantly with overall survival in the Queensland Follow Up cohort (Figure 5C). Particularly, ER^+^/HER2^−^ cases with exclusive nuclear staining (**N1**C0M0) had improved survival (P < 0.0001) while those with strong cytoplasmic staining (Nx**C1**Mx) had poorer survival (P = 0.0066) (Figure 5D). For TNBC, cases with membrane staining in the absence of nuclear staining (N0Cx**M1**) showed significantly poorer survival than the other categories (P = 0.012), accepting the limitation of small numbers in this subset (Figure 5E).

Based on the 389 cases with the associated ER/PR/HER2 IHC subtype and patients survival annotations in the Queensland follow-up cohort, IHC suggests that cases with membrane or strong cytoplasmic KCNN4 expression (Figure 5C: Nx**C1**Mx and N0Cx**M1**, 71 cases, 18.3%), particularly TNBC with membrane expression (Figure 5E: N0Cx**M1**, 10% of TNBC), may benefit from KCNN4 inhibitors, but not cases with exclusive nuclear staining (Figure 5C: **N1**C0M0, 42 cases, 10.6%). These patterns also suggest that KCNN4 has different roles in breast cancer pathology depending on its subcellular localization.

### Dose escalation of Senicapoc

We tested escalating doses of Senicapoc (20, 40, 80 and 120 mg/kg) in non-tumor bearing, immunocompetent BALB/c mice and found that up to 120 mg/kg of Senicapoc injected weekly for 12 weeks was well tolerated without any weight loss (**Supplementary Figure 4**) or other signs of treatment-associated toxicity such as ruffled coat, skin lesion, hunched posture or reluctance to eat or move.

### Senicapoc delays tumor development in a carcinogen-induced mammary carcinoma model

Dimethylbenz[a]anthracene (DMBA) is a chemical carcinogen that can be used to induce ER^+^ tumors in mice [25]. We pre-treated mice with Senicapoc for 6 weeks prior to DMBA exposure and found that it significantly delayed tumor onset compared to vehicle controls; median 22.5 weeks compared to 14 weeks (P = 0.008; Figure 6).

**Figure 6.**
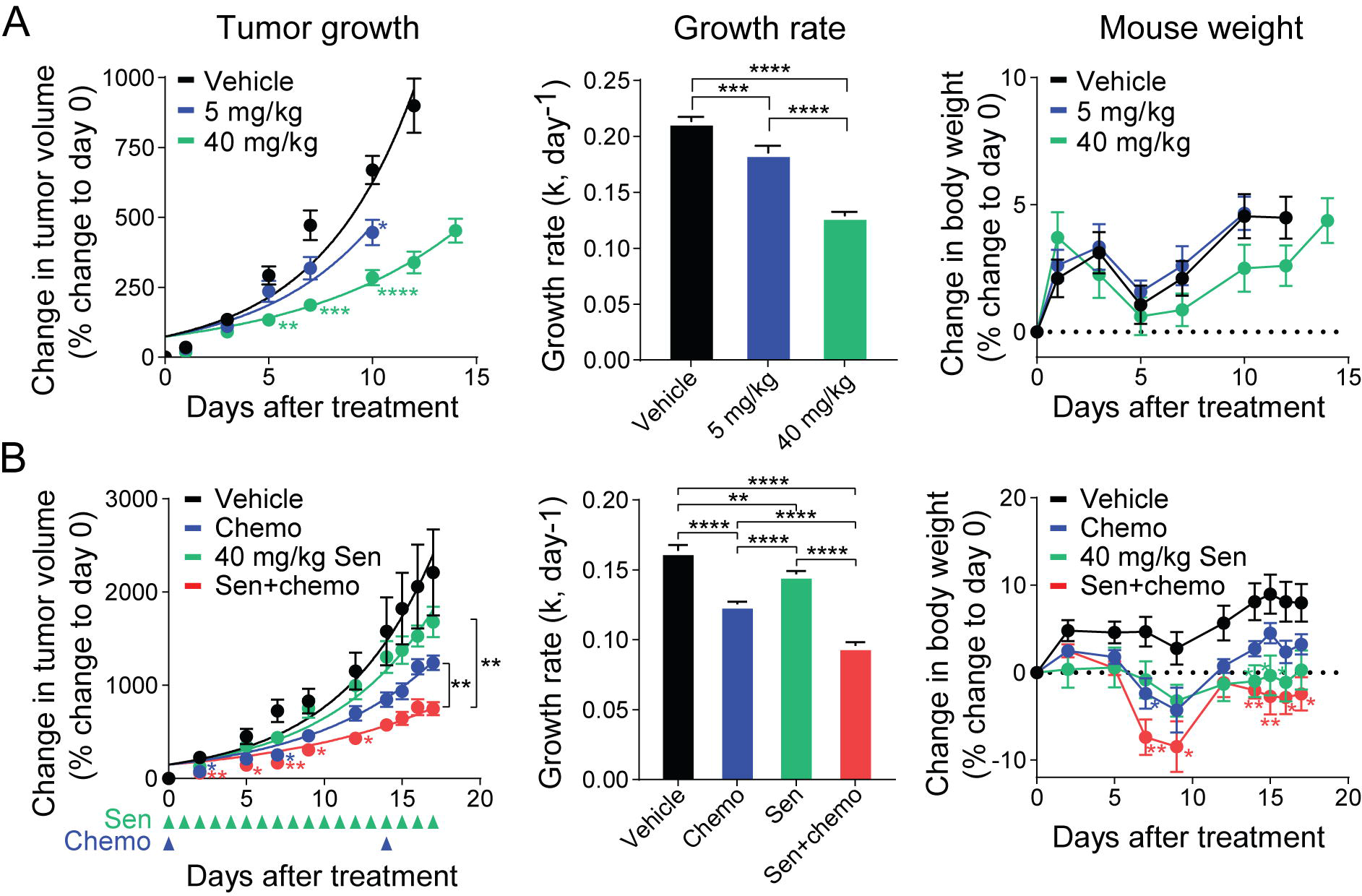
Senicapoc delayed development of DMBA-induced mammary carcinoma. 8-week-old female C57BL/6 mice were treated with 120 mg/kg Senicapoc or vehicle by subcutaneous injection 3 times a week for 6 weeks, then treated with 1 mg DMBA by oral gavage weekly for 6 weeks whilst continuing their Senicapoc dosing. Mammary tumor onset was monitored and latency to 200 mm^3^ is shown (n = 40 per group). Mice that developed lymphomas or gastrointestinal tumors prior to development of mammary tumors were removed from the study.

### Senicapoc delays tumor development in an ER-negative mammary carcinoma model

We assessed the impact of preventative Senicapoc treatment in the SSM1 ER^−^ (Stat1 deficient syngeneic cell line grown in 129SvEv mice. Senicapoc showed a trend toward delayed tumor onset, from 4.8 weeks to 5.35 weeks (P = 0.0588). Senicapoc did significantly slow tumor growth so that survival was increased in Senicapoc-treated mice (median 9.15 weeks compared to 7.3 weeks; P = 0.0064; Figure 7). We found no effect of Senicapoc on the SSM3 ER^−^ (Stat1 deficient) syngeneic cell line (**Supplementary Figure 5**), likely because SSM3 cells express very low levels of *Kcnn4* (Supplementary Figure 6).

**Figure 7.**
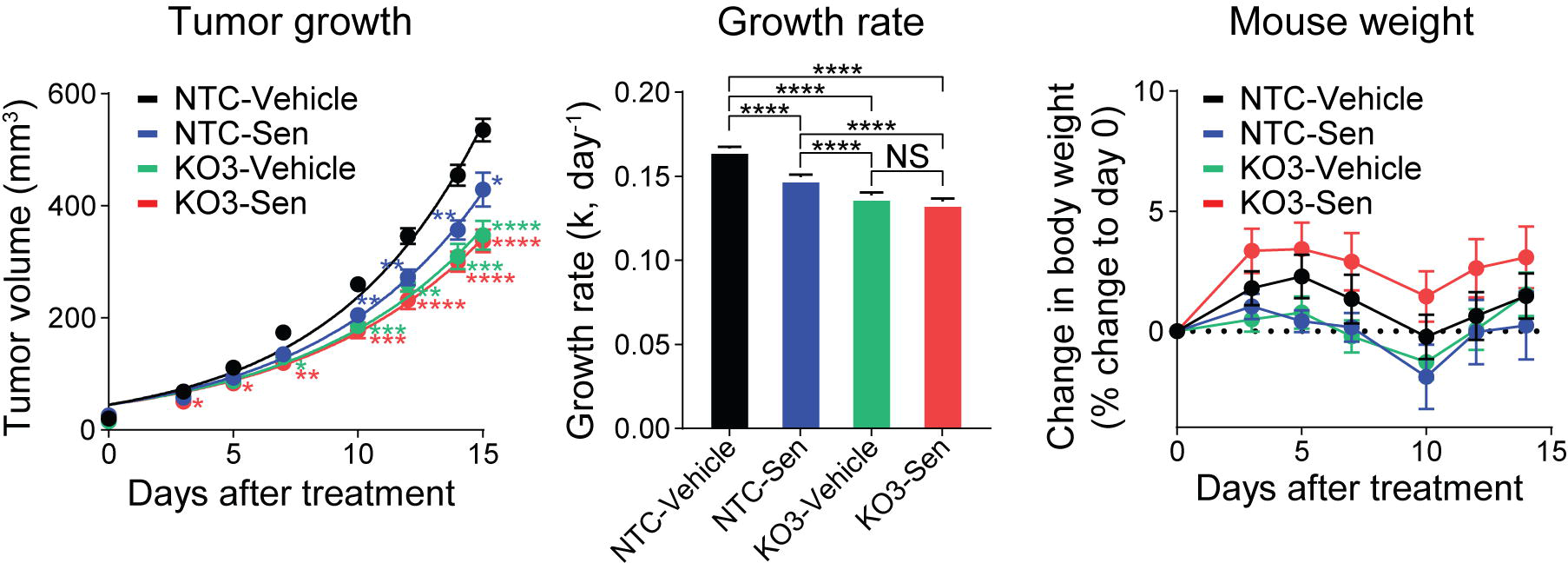
Senicapoc delayed development of ER-negative mammary carcinoma. 8-week-old female 129SvEv mice were treated with 120 mg/kg Senicapoc or vehicle by subcutaneous injection 3 times a week for 6 weeks. They were then injected with 100,000 syngeneic SSM1 cells into the mammary fat pad. The mice were then monitored for mammary tumors while treatment with Senicapoc or vehicle continued. Tumor latency to 200 mm^3^ and survival are shown (n = 12 per group).

### Senicapoc dose-dependent effect on tumor growth in vivo

The 4T1 murine, triple negative, mammary carcinoma cell line is a commonly used model for evaluating mammary tumor growth *in vivo* [30], and expresses KCNN4 (**Supplementary Figure 6**). We evaluated Senicapoc at 2, 20, or 120 mg/kg for anti-tumor effect in BALB/c mice bearing orthotopic murine 4T1 TNBC tumors. At 120 mg/kg injected subcutaneously three times a week, Senicapoc showed a small but a significant delay in tumor growth and improved survival (P = 0.02), without weight loss or other signs of toxicity (Figure 8A).

**Figure 8.** Senicapoc inhibited 4T1 tumor growth *in vivo*. (**A**) Dose-escalation study of Senicapoc in tumor-bearing mice. Immunocompetent BALB/c mice bearing the syngeneic murine breast 4T1 tumors were treated with Senicapoc beginning when tumors reached 50 mm^3^ in volume (day 0). Mice were treated three times a week with 2, 20 or 120 mg/kg Senicapoc (n = 5 per group) which was delivered by subcutaneous injections. (**B**) Senicapoc potentiates chemotherapy efficacy in the 4T1 model. Senicapoc (40 mg/kg) and chemotherapy (docetaxel 2 mg/kg and doxorubicin 10 mg/kg) were used alone or in combination to treat murine 4T1 mammary tumors established in BALB/c mice (n = 10 mice per group). Mice were treated when the tumors reached 25 mm^3^ volume. Drugs were delivered subcutaneously thrice weekly for Senicapoc, and in weeks 1 and 3 for chemotherapy. <u>Left panels</u>: tumor growth curves based on the percentage change of tumor volume compared to day 0 (** P < 0.01, *** P < 0.001, **** P < 0.0001, compared to vehicle group, two-way Anova). <u>Middle panels</u>: tumor growth rates (k, day^−1^) measured from fitting the tumor growth curves to an exponential growth equation using GraphPad Prism (** P < 0.01, **** P < 0.0001, one-way Anova). <u>Right panels</u>: Percentage (%) change in mouse weights compared to initial weights (* P < 0.05, compared to vehicle group, mixed-effects analysis).

### Senicapoc in combination with chemotherapy in vivo

We next combined Senicapoc with doxorubicin and docetaxel, which is a first-line regimen used clinically for TNBC. Weight loss occurred in mice bearing 4T1 tumors whether treated with Senicapoc (120 mg/kg), chemotherapy or both (**Supplementary Figure 7A**). We therefore tested reducing the dose of Senicapoc for the treatment of 4T1 tumors to 80 or 40 mg/kg, administered thrice weekly (**Supplementary Figure 7B**). While the skin lesions remained, along with weight loss in chemotherapy- and combination-treated mice, the lower dose was as efficacious as 80 mg/kg in treating the tumors. Thus, we continued with the reduced dose of Senicapoc (40 mg/kg), and successfully addressed the issue of skin lesions in the 4T1 hosts by trimming their toenails to prevent damage from scratching and by injecting into different locations on the flank (instead of the same location in the neck). We alleviated the weight loss by only administering chemotherapy in weeks 1 and 3 (Figure 8B). Senicapoc monotherapy was as effective as chemotherapy alone in controlling cell growth (P < 0.0001), with the combination treatment better than chemotherapy alone (P < 0.01).

### Senicapoc delivered by oral gavage

Clinically, Senicapoc is administered orally so we tested the efficacy of Senicapoc delivered daily by oral gavage. Using doses that were soluble in the gavage vehicle (5 and 40 mg/kg), Senicapoc significantly inhibited tumor growth in a dose-dependent manner with no weight loss or other signs of adverse events (Figure 9A). In a repeat experiment we found that only three consecutive daily doses of Senicapoc at 40 mg/kg significantly reduced tumor mass (**Supplementary Figure 8**). We next investigated whether Senicapoc retains its combinatorial efficacy with chemotherapy when switched to oral gavage delivery. Notably, the new administration route obviated the need to trim toenails and the treatment maintained efficacious when combined with the original chemotherapy treatments (Figure 9B).

**Figure 9.** Efficacy of Senicapoc delivered by oral gavage. (**A**) Dose-dependent tumor suppression by Senicapoc in the 4T1 model. BALB/c mice bearing syngeneic murine breast 4T1 tumors were treated with Senicapoc beginning when tumors reached 75 mm^3^ on average (day 0). Mice were treated daily with 5 or 40 mg/kg Senicapoc or vehicle control (n = 10 per group) which was delivered by oral gavage. (**B**) Senicapoc given by oral gavage potentiates chemotherapy efficacy in the 4T1 model. Senicapoc (40 mg/kg) and chemotherapy (docetaxel 2 mg/kg and doxorubicin 10 mg/kg) were used alone or in combination to treat murine 4T1 mammary tumors established in BALB/c mice (n = 10 per group). Mice were treated when the tumors reached 30 mm^3^ on average (day 0). Drugs were delivered subcutaneously in weeks 1 and 3 for chemotherapy. <u>Left panels</u>: tumor growth curves based on the percentage change of tumor volume compared to day 0 (* P < 0.05, ** P < 0.01, *** P < 0.001, **** P < 0.0001, compared to vehicle group at different time points or between indicated groups at final time point, two-way Anova/mixed-effects analysis). <u>Middle panels</u>: tumor growth rates (k, day^−1^) measured from fitting the tumor growth curves to an exponential growth equation using GraphPad Prism (** P < 0.01, *** P < 0.001, **** P < 0.0001, one-way Anova). <u>Right panels</u>: Percentage (%) change in mouse weights compared to initial weights (* P < 0.05, ** P < 0.01, compared to vehicle group, mixed-effects analysis).

### KCNN4-knockout to look for off target effects of Senicapoc

In order to determine whether Senicapoc acted solely through inhibition of KCNN4 or has off target effects on other potassium channels, we used the CRISPR/Cas9 system to generate modified 4T1 cells with one non-target control (NTC) gRNA sequence and three gRNAs to generate *Kcnn4* knock out (KO) lines. Out of the three *Kcnn4* KO lines, we selected *Kcnn4* KO3 based on bulk *Kcnn4* mRNA expression (**Supplementary Figure 9**) and made a *Kcnn4* KO3 single cell pool, along with an NTC single cell pool, having validated the Kcnn4 mutations in the single cell clones by ICE analysis of Sanger sequencing traces (**Supplementary Figure 10**). The NTC and *Kcnn4* KO3 lines were pooled from clones with similar *in vitro* growth patterns and they exhibited similar proliferation rates to their parental 4T1 cells *in vitro* in both 2D and 3D models (**Supplementary Figure 11**). However, possibly due to the high Cas9 expression in these cells (**Supplementary Figure 12**), NTC and *Kcnn4* KO3 pooled lines failed to grow tumors in BALB/c mice (**Supplementary Figure 13**). We therefore investigated the off target effects of Senicapoc in BALB/c nude mice. Knockout of *Kcnn4* significantly slowed down tumor growth *in vivo* (Figure 10). Senicapoc treatment by oral gavage consistently retarded NTC tumor growth but did not exert any additional effect in the growth of *Kcnn4* KO3 tumors. These results suggest there was no off-target function of Senicapoc, even though there are other potassium channels expressed in breast cancer [31–33].

**Figure 10:** Effect of Senicapoc on *Kcnn4* KO tumors. BALB/c nude mice bearing 4T1 tumors that were *Kcnn4* KO or wild-type (NTC) were treated with 40 mg/kg Senicapoc or vehicle control by daily oral gavage starting when tumors reached 20 mm^3^ on average (n = 17-18 per group; pooled from two independent experiments). <u>Left panel</u>: tumor growth curves from start of treatment (* P < 0.05, ** P < 0.01, *** P < 0.001, **** P < 0.0001, compared to NTC-vehicle group, two-way Anova). <u>Middle panel</u>: tumor growth rates (k, day^−1^) measured from fitting the tumor growth curves to an exponential growth equation using GraphPad Prism (**** P < 0.0001; one-way Anova). <u>Right</u> <u>panel</u>: Percentage (%) change in mouse weights compared to initial weights.

## Discussion

Genome-wide association studies (GWAS) have identified a locus at 19q13.1 associated with risk of both ER^+^ and ER^−^ breast cancer [34]. Our analysis of public expression data indicates that the target gene at this locus is *KCNN4*, and that the allele associated with increased risk is associated with increased expression of *KCNN4*. This suggests that a KCNN4 inhibitor, Senicapoc, may have potential for preventing breast cancer. The only approved risk reduction medication for breast cancer available for use in high risk women is Tamoxifen, which targets another risk-associated gene, *ESR1* [35] but uptake is low in clinical practice in part because of concerns about the side effects [36–39]. We therefore tested Senicapoc in two models of murine mammary carcinoma, and found that it delayed tumor development.

As Tamoxifen is effective not only as a preventative, but also a therapeutic strategy for ER^+^ breast cancer, we also considered whether Senicapoc could decrease tumor growth when used as a monotherapy, or in combination with chemotherapy. First we evaluated the expression of KCNN4, Senicapoc’s target, in breast cancer. We found that in the TCGA cohort, *KCNN4* expression was significantly higher in TNBC and BLBC cases compared to all other subtypes, suggesting that inhibition of KCNN4 might be especially useful for the treatment of these subtypes. We then extended our analysis to look at protein expression and subcellular localization in a breast cancer patient cohort with up to 25 years of follow up. We found that nuclear KCNN4 expression was strongly associated with ER positivity. Exclusive nuclear expression, which was observed in 18% of ER^+^/HER2^−^ breast cancer, predicted better survival in this subtype. Cytoplasmic KCNN4 expression was observed in 11% and 15% of the ER^+^/HER2^−^ and TNBC subtypes, respectively, and predicted poorer survival in ER^+^/HER2^−^ breast cancer. However, membrane KCNN4 expression was mainly restricted to TNBC, with the 10% of TNBC with membrane staining having dismal survival. While we do not have an explanation for the nuclear and cytoplasmic subcellular localization of KCNN4, this has been observed previously [40], and warrants further investigation to determine the function of KCNN4 in these contexts.

We focused our studies of KCNN4 inhibition on 4T1 cells, a mouse model of TNBC, because of the paucity of targeted therapies for this aggressive tumor subtype. We found that Senicapoc alone can reduce the growth of 4T1 tumors, as effectively as chemotherapy, and that the combination of Senicapoc and chemotherapy is even more effective. By making a *KCNN4* knockout derivative of 4T1, we showed that although this did not affect the growth of the cells *in vitro*, it had a similar effect on tumor growth *in vivo* as the use of Senicapoc. This suggests that Senicapoc is acting on Kcnn4 and has no off-target effects. Senicapoc had not previously been tested in mouse models of TNBC, and further experiments are warranted in other syngeneic models of mammary carcinoma.

Given that it was well tolerated in clinical trials for other diseases, our data suggest that clinical testing of Senicapoc is warranted for the treatment of TNBC, and reduction of risk. Not only is Senicapoc orally bioavailable, but it also has much longer half-life in humans than in rodents [7], suggesting that even lower doses might be efficacious to prevent or treat breast cancer. The human equivalent dose for 40 mg/kg in mice is 3.3 mg/kg, but the adult human clinical trial for sickle cell anemia used only a single 150 mg Senicapoc loading dose followed by 10 mg daily maintenance dose [17], and the clinical trial for asthma used 40 mg/day [17].

Although we focused our pre-clinical studies mostly on TNBC, other subtypes of breast cancer might also respond to Senicapoc. Examples include cases where high KCNN4 expression levels are associated with a worse outcome, such as ER^+^/HER2^−^ breast cancer with strong cytoplasmic expression, and recurrent tumors that still express KCNN4 after chemotherapy. In this context, it will be important to build prospective IHC assessment of KCNN4 into any forthcoming adjuvant or neoadjuvant clinical trials to see whether this could be developed as a companion biomarker.

## Conclusion

Our study found that *KCNN4* is associated with chemotherapy resistance in breast cancer, and the membrane and cytoplasmic localization of its protein product is associated with aggressive clinicopathological features and poor prognosis. Our *in vivo* studies demonstrated that the well-tolerated KCNN4 inhibitor, Senicapoc, may be useful for prevention of breast cancer in some situations, and has anti-tumor activity in relevant preclinical models and suggested that combining it with chemotherapy may provide added benefit to patients with TNBC.

## Abbreviations

GWAS: Genome-wide association studies
CCV: candidate causal variant
ER: estrogen receptor
TNBC: triple negative breast cancer
PR: progesterone receptor
HER2: human epidermal growth factor receptor 2
eQTL: expression quantitative trait loci
GTEx: Genotype-Tissue Expression
RT-qPCR: quantitative real time polymerase chain reaction
IHC: Immunohistochemistry
DMBA: Dimethylbenz[a]anthracene
NTC: non-target control
TCGA: The Cancer Genome Atlas
BLBC: basal-like breast cancer
PAM: Prediction Analysis of Microarray.

## Declarations

### Ethics approval and consent to participate

The use of clinical samples and patient data for this study was approved by human research ethics committees at The University of Queensland (ref. 2005000785) and the Royal Brisbane and Women’s Hospital (2005/022). All animal work was conducted in accordance with the National Health and Medical Research Council guidelines under the approval of the QIMR Berghofer or Peter MacCallum Cancer Centre Animal Ethics Committees.

### Consent for publication

Not applicable.

### Availability of data and material

All data generated or analyzed during this study are included in this published article and its supplementary materials, or in indicated references. Supplementary materials include two excel files and thirteen supplementary Figures. The excel files include sequences of primers, transcripts, shRNAs and sgRNAs used in the experiments, and the data on the association of KCNN4 localization with clinicopathological variables. The supplementary Figures include data that support the main Figures in the manuscript, including patient survival curves based on KCNN4 expression, *KCNN4* and *Kcnn4* expression in mammary cell lines, data on KCNN4 antibody validation, as well as safety data, *in vivo* efficacy and optimization using TNBC models.

### Authors’ contributions

CX, JB, GRM, GCT, KLB and FA conceptualized and supervised the study.

KLB, GCT, JB and FA provided resources/obtained funding.

CX, MM, WS, GCT, KLB, KM and FA designed the methodology

CX, MM, WS, DMB, AC, ZT, HYH, XC, MP, MR, SJ, DCC, JRK, JMS, AEM, LK, FA and DB performed the investigations/experiments.

JMS and SRL generated the breast cancer cohort.

FA carried out the in silico analysis/data curation.

KLB, GCT, CX writing reviewing and editing

CX writing the original draft

## Acknowledgements

We thank the patients who donated samples and the Brisbane Breast Bank for collection, annotation and provision of clinical samples for IHC studies. We thank Dr Robert D. Schreiber for SSM1 and SSM3 cell lines. We acknowledge the support of Metro North Hospital & Health Service in relation to collection of clinical subject data and materials, and thank the QIMR Berghofer the Animal, Flow, Microscopy and Histology facilities for technical support.

## Supplementary Figure Legends

**Supplementary Figure 1. High *KCNN4* does not associate with survival after endocrine therapy and radiotherapy.** Overall survival (OS), relapse-free survival (RFS) or distant metastasis-free survival (DMFS) of TCGA breast cancer cases were compared after categorizing cases by treatment regimen and *KCNN4* expression quartiles (Q1: bottom; Q2/Q3: middle quartiles; Q4: top quartile). Log-rank test was used for statistical analysis.

**Supplementary Figure 2. *KCNN4* expression in human breast tumor cell lines.** Expression of *KCNN4* in human breast cancer cell lines was measured by RT-qPCR. Three primer sets were used for *KCNN4* (refer to Supplementary Table 1 for primers and transcripts details). Relative expression (2^−ΔΔCt^) is shown (n = 3, error bars are SEM), using housekeeping genes as references and normalized to the expression in T47D.

**Supplementary Figure 3. Validation and vasculature signatures of the staining of the antibody against human KCNN4.** (**A** and **B**) MDA-MB-231 cells expressing non-target control shRNA (shNTC) or shRNAs against *KCNN4* (shRNA#3 and shRNA#5) were analyzed by RT-qPCR (**A**) and immunofluorescence and IHC (**B**) to confirm specificity of the antibody (Sigma, clone HPA053841). *** P < 0.001, **** P < 0.0001, compared to shNTC of each primer set, one-way ANOVA. (**C**) Representative IHC images showing patterns of KCNN4 staining in tumor-associated vasculature.

**Supplementary Figure 4. Dose-escalation study of Senicapoc in non-tumor bearing mice.** Non-tumor bearing, wild-type BALB/c mice received escalating doses of Senicapoc starting from 8-weeks of age. Senicapoc, and vehicle control, was administered weekly by subcutaneous injection for 12 weeks. Mice were monitored for weight loss and data shown is the % change in mouse weight compared to initial weight (n = 5 mice per group). Senicapoc was well tolerated without weight loss or other signs of treatment-associated toxicity such as ruffled coat, skin lesion, hunched posture or reluctance to eat or move.

**Supplementary Figure 5. Senicapoc did not delay development of ER-negative mammary carcinoma SSM3.** 8-week-old female 129SvEv mice were treated with 120 mg/kg Senicapoc or vehicle by subcutaneous injection 3 times a week for 6 weeks. They were then injected with 100,000 syngeneic SSM3 cells into the mammary fat pad. The mice were then monitored for mammary tumors while treatment with Senicapoc or vehicle continued. Tumor latency to 200 mm^3^ and tumor growth are shown (n = 12 per group).

**Supplementary Figure 6. *Kcnn4* expression in murine mammary tumor lines.** Expression of *Kcnn4* in murine mammary tumor cell lines was measured by RT-qPCR. Four primer sets were used for *Kcnn4* (refer to Supplementary Table 1 for primers and transcripts details). Relative expression (2^−ΔΔCt^) is shown (n = 3, error bars are SEM), using housekeeping genes as references and normalized to the expression in SSM3. ** P < 0.01, *** P < 0.001, unpaired t-test.

**Supplementary Figure 7. Optimization of Senicapoc treatment in combination with chemotherapy.** (**A**) High dose Senicapoc in combination with chemotherapy. Senicapoc (120 mg/kg) and chemotherapy (docetaxel 2 mg/kg and doxorubicin 10 mg/kg) were used alone or in combination to treat murine 4T1 mammary tumors established in BALB/c mice (n = 7 mice per group). The mice were treated when tumors reached 50 mm^3^ in volume and drugs were delivered subcutaneously every two days for Senicapoc, but chemotherapy was only administered in week 1 because of weight loss. (**B**) Senicapoc dose reduction for combination with chemotherapy. Senicapoc (80 or 40 mg/kg) and chemotherapy (docetaxel 2 mg/kg and doxorubicin 10 mg/kg) were used alone or in combination to treat murine 4T1 mammary tumors established in BALB/c mice (n = 6-12 mice per group). The mice were treated when tumors reached 50 mm^3^ in volume and drugs were delivered subcutaneously three times a week for Senicapoc, but weekly for chemotherapy. <u>Left panels:</u> tumor growth curves based on the percentage change of tumor volume compared to day 0 (* P < 0.05, ** P < 0.01, *** P < 0.001, **** P < 0.0001, compared to vehicle group, two-way Anova). <u>Middle panels</u>: tumor growth rates (k, day^−1^) measured from fitting the tumor growth curves to an exponential growth equation using GraphPad Prism (**** P < 0.0001, one-way Anova). <u>Right panels</u>: Percentage (%) change in mouse weights compared to initial weights on day 0.

**Supplementary Figure 8. Senicapoc given by oral gavage significantly reduced tumor growth after three doses.** BALB/c mice bearing mammary 4T1 tumors were treated with 40 mg/kg Senicapoc or vehicle control by daily oral gavage starting when tumors reached 30 mm^3^ on average (n = 4-5 per group). Tumors were sampled three days after first treatment and weighed on a scale. * P < 0.05, student’s t-test.

**Supplementary Figure 9. *Kcnn4* expression in CRISPR-modified 4T1 cells.** Three derivatives of 4T1 cells with *Kcnn4* knockout (KO1, 2 and 3), along with a non-target control (NTC) were generated with CRISPR/Cas9 system. *Kcnn4* mRNA was determined by RT-qPCR at passages 3, 7, and 17 after antibiotic selection. Relative expression (2^−ΔΔCt^) is calculated with the average of the primer sets, using housekeeping genes as references and normalized to the expression of NTC cells. * P < 0.05, ** P < 0.01, *** P < 0.001, **** P < 0.0001, compared to NTC, two-way ANOVA.

**Supplementary Figure 10. Validation of wild-type and knocked-out *Kcnn4* in NTC and KO3 single clones by sequencing.** For each of the single clones selected for NTC and KO3 pools, genomic DNA was extracted for sequencing and Synthego ICE analysis was performed.

**Supplementary** Figure 11**. *In vitro* proliferation of CRISPR-modified 4T1 cells.** For the 2D proliferation assay, cells were cultured over a 72 hour period, with an initial concentration of 20,000 cells per well of 24-well plates. For the 4T1 parental control, 4T1 NTC pre-sort and 4T1 *Kcnn4* KO3 pre-sort cells, 4 wells in each plate were used, and for the 4T1 NTC and *Kcnn4* KO3 single clone pools 6 wells in each plate were used. At the end of incubation, a CellTiter Glo assay was performed and the luminescence signal for each well was recorded. For the 3D proliferation assay, a growth in low attachment (GILA) assay was performed. Using the same 5 cell lines, 10,000 cells per well were plated in two 24-well low attachment plates. Sample sizes were the same as 2D assay. The colonies were allowed to grow for 72 hours in a humidified incubator. At the end of this period a CellTiter Glo assay was performed and the luminescence signal for each well was recorded. Results shown were normalized to the level of parental 4T1 cells.

**Supplementary Figure 12. *Cas9* expression in the single clones that were used to generate NTC and KO3 pools.** *Cas9* mRNA was determined by RT-qPCR in duplicates and relative expression (2^−ΔΔCt^) is calculated with the average of the two primer sets, using housekeeping genes as references and normalized to the expression of parental 4T1 cells.

**Supplementary Figure 13. *In vivo* growth of NTC and *Kcnn4* KO3 pools in BALB/c mice.** 4T1 NTC or *Kcnn4* KO3 pool or parental 4T1 cells prepared in the same way were injected into the mammary fat pad of 8-week old female BALB/c mice and tumor volume was recorded (n = 5). <u>Left</u> <u>panel:</u> tumor growth curves. <u>Right panel</u>: tumor growth rates (k, day^−1^) measured from fitting the tumor growth curves to an exponential growth equation using GraphPad Prism.

## References

1. Michailidou K, Hall P, Gonzalez-Neira A, Ghoussaini M, Dennis J, Milne RL, et al. Large-scale genotyping identifies 41 new loci associated with breast cancer risk. Nat Genet. 2013;45:353–61, 361e1–2.

2. Fachal L, Aschard H, Beesley J, Barnes DR, Allen J, Kar S, et al. Fine-mapping of 150 breast cancer risk regions identifies 191 likely target genes. Nat Genet. 2020;52:56–73.

3. Nelson MR, Tipney H, Painter JL, Shen J, Nicoletti P, Shen Y, et al. The support of human genetic evidence for approved drug indications. Nat Genet. 2015;47:856–60.

4. Suen JC, Hatheway CL, Steigerwalt AG, Brenner DJ. Genetic confirmation of identities of neurotoxigenic Clostridium baratii and Clostridium butyricum implicated as agents of infant botulism. J Clin Microbiol. 1988;26:2191–2.

5. King EA, Davis JW, Degner JF. Are drug targets with genetic support twice as likely to be approved? Revised estimates of the impact of genetic support for drug mechanisms on the probability of drug approval. PLoS Genet. 2019;15:e1008489.

6. Ochoa D, Karim M, Ghoussaini M, Hulcoop DG, McDonagh EM, Dunham I. Human genetics evidence supports two-thirds of the 2021 FDA-approved drugs. Nat Rev Drug Discov. 2022;21:551.

7. Brown BM, Pressley B, Wulff H. KCa3.1 Channel Modulators as Potential Therapeutic Compounds for Glioblastoma. Curr Neuropharmacol. 2018;16:618–26.

8. Jensen BS, Strøbaek D, Olesen SP, Christophersen P. The Ca2+-activated K+ channel of intermediate conductance: a molecular target for novel treatments? Curr Drug Targets. 2001;2:401– 22.

9. Faouzi M, Hague F, Geerts D, Ay A-S, Potier-Cartereau M, Ahidouch A, et al. Functional cooperation between KCa3.1 and TRPC1 channels in human breast cancer: Role in cell proliferation and patient prognosis. Oncotarget. 2016;7:36419–35.

10. Lin P, Li J, Ye F, Fu W, Hu X, Shao Z, et al. KCNN4 induces multiple chemoresistance in breast cancer by regulating BCL2A1. Am J Cancer Res. 2020;10:3302–15.

11. Du Y, Song W, Chen J, Chen H, Xuan Z, Zhao L, et al. The potassium channel KCa3.1 promotes cell proliferation by activating SKP2 and metastasis through the EMT pathway in hepatocellular carcinoma. Int J Cancer [Internet]. 2019; Available from: http://dx.doi.org/10.1002/ijc.32121

12. Jäger H, Dreker T, Buck A, Giehl K, Gress T, Grissmer S. Blockage of intermediate-conductance Ca2+-activated K+ channels inhibit human pancreatic cancer cell growth in vitro. Mol Pharmacol. 2004;65:630–8.

13. Parihar AS, Coghlan MJ, Gopalakrishnan M, Shieh C-C. Effects of intermediate-conductance Ca2+-activated K+ channel modulators on human prostate cancer cell proliferation. Eur J Pharmacol. 2003;471:157–64.

14. Grössinger EM, Weiss L, Zierler S, Rebhandl S, Krenn PW, Hinterseer E, et al. Targeting proliferation of chronic lymphocytic leukemia (CLL) cells through KCa3.1 blockade. Leukemia. Nature Publishing Group; 2014;28:954–8.

15. Zhang P, Yang X, Yin Q, Yi J, Shen W, Zhao L, et al. Inhibition of SK4 Potassium Channels Suppresses Cell Proliferation, Migration and the Epithelial-Mesenchymal Transition in Triple-Negative Breast Cancer Cells. PLoS One. 2016;11:e0154471.

16. Steudel FA, Mohr CJ, Stegen B, Nguyen HY, Barnert A, Steinle M, et al. SK4 channels modulate Ca2+ signalling and cell cycle progression in murine breast cancer. Mol Oncol. 2017;11:1172–88.

17. Ataga KI, Smith WR, De Castro LM, Swerdlow P, Saunthararajah Y, Castro O, et al. Efficacy and safety of the Gardos channel blocker, senicapoc (ICA-17043), in patients with sickle cell anemia. Blood. 2008;111:3991–7.

18. Wulff H, Castle NA. Therapeutic potential of KCa3.1 blockers: recent advances and promising trends. Expert Rev Clin Pharmacol. 2010;3:385–96.

19. Foley CN, Staley JR, Breen PG, Sun BB, Kirk PDW, Burgess S, et al. A fast and efficient colocalization algorithm for identifying shared genetic risk factors across multiple traits. Nat Commun. 2021;12:764.

20. Chan SR, Vermi W, Luo J, Lucini L, Rickert C, Fowler AM, et al. STAT1-deficient mice spontaneously develop estrogen receptor α-positive luminal mammary carcinomas. Breast Cancer Res. 2012;14:R16.

21. Fowler AM, Chan SR, Sharp TL, Fettig NM, Zhou D, Dence CS, et al. Small-animal PET of steroid hormone receptors predicts tumor response to endocrine therapy using a preclinical model of breast cancer. J Nucl Med. 2012;53:1119–26.

22. Schindelin J, Arganda-Carreras I, Frise E, Kaynig V, Longair M, Pietzsch T, et al. Fiji: an open-source platform for biological-image analysis. Nat Methods. 2012;9:676–82.

23. Al-Ejeh F, Simpson PT, Sanus JM, Klein K, Kalimutho M, Shi W, et al. Meta-analysis of the global gene expression profile of triple-negative breast cancer identifies genes for the prognostication and treatment of aggressive breast cancer. Oncogenesis. 2014;3:e100.

24. Raghavendra A, Kalita-de Croft P, Vargas AC, Smart CE, Simpson PT, Saunus JM, et al. Expression of MAGE-A and NY-ESO-1 cancer/testis antigens is enriched in triple-negative invasive breast cancers. Histopathology. 2018;73:68–80.

25. Abba MC, Zhong Y, Lee J, Kil H, Lu Y, Takata Y, et al. DMBA induced mouse mammary tumors display high incidence of activating Pik3caH1047 and loss of function Pten mutations. Oncotarget. 2016;7:64289–99.

26. Stocker JW, De Franceschi L, McNaughton-Smith GA, Corrocher R, Beuzard Y, Brugnara C. ICA-17043, a novel Gardos channel blocker, prevents sickled red blood cell dehydration in vitro and in vivo in SAD mice. Blood. 2003;101:2412–8.

27. Balko JM, Cook RS, Vaught DB, Kuba MG, Miller TW, Bhola NE, et al. Profiling of residual breast cancers after neoadjuvant chemotherapy identifies DUSP4 deficiency as a mechanism of drug resistance. Nat Med. 2012;18:1052–9.

28. Guarneri V, Piacentini F, Ficarra G, Frassoldati A, D’Amico R, Giovannelli S, et al. A prognostic model based on nodal status and Ki-67 predicts the risk of recurrence and death in breast cancer patients with residual disease after preoperative chemotherapy. Ann Oncol. 2009;20:1193–8.

29. Uhlén M, Fagerberg L, Hallström BM, Lindskog C, Oksvold P, Mardinoglu A, et al. Proteomics. Tissue-based map of the human proteome. Science. 2015;347:1260419.

30. Heppner GH, Miller FR, Shekhar PM. Nontransgenic models of breast cancer. Breast Cancer Res. 2000;2:331–4.

31. Liu G-X, Yu Y-C, He X-P, Ren S-N, Fang X-D, Liu F, et al. Expression of eag1 channel associated with the aggressive clinicopathological features and subtype of breast cancer. Int J Clin Exp Pathol. 2015;8:15093–9.

32. Dookeran KA, Zhang W, Stayner L, Argos M. Associations of two-pore domain potassium channels and triple negative breast cancer subtype in The Cancer Genome Atlas: systematic evaluation of gene expression and methylation. BMC Res Notes. 2017;10:475.

33. Oeggerli M, Tian Y, Ruiz C, Wijker B, Sauter G, Obermann E, et al. Role of KCNMA1 in breast cancer. PLoS One. 2012;7:e41664.

34. Michailidou K, Lindström S, Dennis J, Beesley J, Hui S, Kar S, et al. Association analysis identifies 65 new breast cancer risk loci. Nature. 2017;551:92–4.

35. Dunning AM, Michailidou K, Kuchenbaecker KB, Thompson D, French JD, Beesley J, et al. Breast cancer risk variants at 6q25 display different phenotype associations and regulate ESR1, RMND1 and CCDC170. Nat Genet. 2016;48:374–86.

36. Meiser B, Wong WKT, Peate M, Julian-Reynier C, Kirk J, Mitchell G. Motivators and barriers of tamoxifen use as risk-reducing medication amongst women at increased breast cancer risk: a systematic literature review. Hered Cancer Clin Pract. 2017;15:14.

37. Smith SG, Sestak I, Forster A, Partridge A, Side L, Wolf MS, et al. Factors affecting uptake and adherence to breast cancer chemoprevention: a systematic review and meta-analysis. Ann Oncol. 2016;27:575–90.

38. Donnelly LS, Evans DG, Wiseman J, Fox J, Greenhalgh R, Affen J, et al. Uptake of tamoxifen in consecutive premenopausal women under surveillance in a high-risk breast cancer clinic. Br J Cancer. 2014;110:1681–7.

39. Bober SL, Hoke LA, Duda RB, Regan MM, Tung NM. Decision-making about tamoxifen in women at high risk for breast cancer: clinical and psychological factors. J Clin Oncol. 2004;22:4951–7.

40. Zhao H, Guo E, Hu T, Sun Q, Wu J, Lin X, et al. KCNN4 and S100A14 act as predictors of recurrence in optimally debulked patients with serous ovarian cancer. Oncotarget. 2016;7:43924– 38.

