## Supplementary material for "The potential of Senicapoc, a KCNN4 inhibitor, for the prevention and treatment of breast cancer": Supple Figures

**Supplementary Table 1: qRT-PCR and DNA sequencing primer sets, human KCNN4 transcripts, human KCNN4 shRNAs, and sgRNA sequences**

| Human Primers | sequence | Length | Tm | Location |
|---|---|---|---|---|
| **KCNN4-1F** | TTGGCTGATCCCCATCACATT | 21 | 61.5 | 723-743 |
| **KCNN4-1R** | CAGGCTTCTTGTAGCACTCGG | 21 | 62.7 | 968-948 |
| **KCNN4-2F** | CTGCTGCGTCTCTACCTGG | 19 | 61.8 | 469-487 |
| **KCNN4-2R** | AGGGTGCGTGTTCATGTAAAG | 21 | 60.6 | 612-592 |
| **KCNN4-3R** | AAGCTCCGGGAACAAGTGAAC | 21 | 62.6 | 1078-1098 |
| **KCNN4-3F** | CGCCAGCGTGTCAATCTGT | 19 | 62.9 | 1200-1182 |

| Mouse Primers | sequence |
|---|---|
| **mKcnn4-1F** | CTGATTCCGATCACATTCCTGAC |
| **mKcnn4-1R** | TTTCTCCGCCTTGTTGAACTC |
| **mKcnn4-2F** | TTCAACAAGGCGGAGAAACAC |
| **mKcnn4-2R** | TCTTGCGCTGATGTCTGCG |
| **mKcnn4-3F** | TACGGCTGAAACACCGGAAG |
| **mKcnn4-3R** | TGCCAGACCGTCGATTCTCT |
| **mKcnn4-4F** | GCTCAACCAAGTCCGCTTC |
| **mKcnn4-4R** | GTGATCGGAATCAGCCACAGT |

| House keeping genes | Primer sequence |
|---|---|
| **MS_Gapdh_FWD** | **AGGTCGGTGTGAACGGATTTG** |
| **MS_Gapdh_REV** | **TGTAGACCATGTAGTTGAGGTCA** |
| **MS_Actb_FWD** | **GGCTGTATTCCCCTCCATCG** |
| **MS_Actb_REV** | **CCAGTTGGTAACAATGCCATGT** |
| hu_ACTIN_qPCR_Fw | CCAAGGCCAACCGCGAGAAGATGAC |
| hu_ACTIN_qPCR_Rv | AGGGTACATGGTGGTGCCGCCAGAC |
| hu_18S rRN_Fw | AACTTTCGATGGTAGTCGCCG |
| hu_18S rRNA_Rv | CCTTGGATGTGGTAGCCGTTT |

| human transcripts | Transcript ID | bp | Protein | Biotype | UniProt | Primer Set 1 | Primer Set 2 | Primer Set 3 |
|---|---|---|---|---|---|---|---|---|
| **KCNN4-210** | ENST00000648319.1 | 1956 | 427aa | Protein coding | O15554 | **Yes** | **Yes** | **Yes** |
| **KCNN4-208** | ENST00000615047.4 | 2029 | 295aa | Protein coding | D1MQ08 | **Yes** | | **Yes** |
| **KCNN4-206** | ENST00000600909.1 | 1190 | 59aa | Protein coding | M0QZ70 | | | |
| **KCNN4-202** | ENST00000598836.1 | 1003 | 165aa | Protein coding | M0R1J0 | | | **Yes** |
| **KCNN4-204** | ENST00000599720.5 | 983 | 57aa | Nonsense mediated decay | M0R2F1 | **Yes** | | **Yes** |
| **KCNN4-205** | ENST00000600408.1 | 648 | 113aa | Nonsense mediated decay | M0R2E8 | **Yes** | | **Yes** |
| **KCNN4-207** | ENST00000601549.2 | 1406 | No protein | Retained intron | - | **Yes** | **Yes** | **Yes** |
| **KCNN4-203** | ENST00000599107.1 | 687 | No protein | Retained intron | - | | **Yes** | |
| **KCNN4-201** | ENST00000597184.5 | 631 | No protein | Retained intron | - | | | **Yes** |
| **KCNN4-209** | ENST00000648053.1 | 1270 | No protein | lncRNA | - | **Yes** | | **Yes** |

| | | | |
|---|---|---|---|
| shRNA 1 | TRCN0000301090 | CCGGGCCTGGATGTTCTACAAACATCTCGAGATGTTTGTAGAACATCCAGGCTTTTTG | 2 |
| shRNA 2 | TRCN0000301089 | CCGGCAAGATGCACATGATCCTGTACTCGAGTACAGGATCATGTGCATCTTGTTTTTG | 2 |
| shRNA 3 | TRCN0000323045 | CCGGCATGATGGATATCCAGTATACCTCGAGGTATACTGGATATCCATCATGTTTTTG | 2 |
| shRNA 4 | TRCN0000043933 | CCGGGCCTGGATGTTCTACAAACATCTCGAGATGTTTGTAGAACATCCAGGCTTTTTG | 1 |
| shRNA 5 | TRCN0000043935 | CCGGCAAGATGCACATGATCCTGTACTCGAGTACAGGATCATGTGCATCTTGTTTTTG | 1 |
| NTC | SHC016 | CCGGCAACAAGATGAAGAGCACCAACTCGAGTTGGTGCTCTTCATCTTGTTGTTTTT | 1 |
| **shRNA ID*** | **Clone ID** | **Target Sequence** | **TRC version** |

* **Note**:Sequences of shRNA1 and shRNA4, and of shRNA2 and shRNA5, are identical but with different TRC versions. The TRC2 vector has an added WPRE (or the Woodchuck Hepatitis Post-Transcriptional Regulatory Element)

| | sgRNA Sequence |
|---|---|
| sgRFP | GATGCCCGGCTTCCACTTCG |
| mKcnn4_sg1 | GGCAGGCTGTCAATGCCACG |
| mKcnn4_sg2 | CTGTACATGAACACGCACCC |
| mKcnn4_sg3 | GCAGACAATCTTGCCCCACA |

| DNA sequencing primers for murine K | Sequence |
|---|---|
| mKC1 Seq_FWD1 | TCCTCCAGGCTTGACCAGACCT |
| mKC1 Seq_REV1 | TGGCCCATGGTTTCCTGAGCCA |
| mKC2 Seq_FWD1 | AAGGCGAGGCGTTGCTGTCC |
| mKC2 Seq_REV1 | GGCTCAGAAGTTGGGAATCACCTCAA |
| mKC2 Seq_FWD2 | GTTGCTGTCCCTGGCCATGCTC |
| mKC2 Seq_REV2 | TCTGGGTGGCTCAGAAGTTGGGA |
| mKC3 Seq_FWD1 | GGCTGGAAGAGGAACAGTATGGGGG |
| mKC3 Seq_REV1 | CTCCCAAGTGCCCGGCAGATCT |

**Supplementary Table 2: Association of KCNN4 subcellular localisation with clinicopathological variables.**

| *P-value calculated using Chi-square test. ~ Fisher's exact test. * Chi-square test for trend* | | total | Membrane KCNN4 staining n (%) | | | Cytoplasmic KCNN4 staining n (%) | | | | Nuclear KCNN4 staining n (%) | | | Consolidated KCNN4 staining patterns | | | | | Vasculature KCNN4 staining n (%) | | |
|---|---|---|---|---|---|---|---|---|---|---|---|---|---|---|---|---|---|---|---|---|
| | | | Positive (M1) | Negative (M0) | *p* value~ | Strong (C2) | Weak (C1) | Negative (C0) | *p* value~ | Positive (N1) | Negative (N0) | *p* value~ | n (%) N0CxM1 | n (%) NxC1Mx | n (%) N1C0M0 | n (%) N0C0M0 | *p* value~ | Positive | Negative | *p* value~ |
| Histological type | ductal NOS | 258 | 6 (2.3) | 252 (97.7) | | 33 (12.8) | 64 (24.8) | 161 (62.4) | | 53 (20.5) | 205 (79.5) | | 5 (71.4) | 30 (66.7) | 49 (73) | 174 (60.2) | | 168 (65.1) | 90 (34.9) | |
| | lobular/variants | 45 | 0 (0) | 45 (100) | | 3 (6.7) | 7 (15.6) | 35 (77.8) | | 6 (13.3) | 39 (86.7) | | 0 (0) | 3 (6.7) | 5 (7.5) | 37 (12.8) | | 28 (62.2) | 17 (37.8) | |
| | mixed ducto-lob | 28 | 0 (0) | 28 (100) | | 3 (10.7) | 9 (32.1) | 16 (57.1) | | 4 (14.3) | 24 (85.7) | | 0 (0) | 3 (6.7) | 4 (6) | 21 (7.3) | | 19 (67.9) | 9 (32.1) | |
| | mixed | 34 | 2 (5.9) | 32 (94.1) | 0.3898 | 5 (14.7) | 8 (23.5) | 21 (61.8) | 0.4474 | 3 (8.8) | 31 (91.2) | 0.4787 | 2 (28.6) | 3 (6.7) | 3 (4.5) | 26 (9) | 0.6177 | 25 (73.5) | 9 (26.5) | 0.9173 |
| | metaplastic | 14 | 0 (0) | 14 (100) | | 3 (21.4) | 1 (7.1) | 10 (71.5) | | 2 (14.3) | 12 (85.7) | | 0 (0.0) | 3 (6.7) | 2 (3) | 9 (3.1) | | 10 (71.4) | 4 (28.6) | |
| | special types | 29 | 0 (0) | 29 (100) | | 3 (10.3) | 4 (13.8) | 22 (75.9) | | 4 (13.8) | 25 (86.2) | | 0 (0.0) | 3 (6.7) | 4 (6) | 22 (7.6) | | 19 (65.5) | 10 (34.5) | |
| | *n* | *408* | *8 (2)* | *400 (98)* | | *50 (12.3)* | *93 (22.7)* | *265 (65)* | | *72 (17.6)* | *336 (82.4)* | | *7 (1.7)* | *45 (11)* | *67 (16.4)* | *289 (70.9)* | | *269 (65.9)* | *139 (34.1)* | |
| Grade | 1 | 57 | 0 (0) | 57 (100) | | 2 (3.5) | 9 (15.8) | 46 (80.7) | | 15 (26.3) | 42 (73.7) | | 0 (0) | 2 (4.4) | 15 (19.5) | 40 (13.8) | | 39 (68.4) | 18 (31.6) | |
| | 2 | 200 | 2 (1) | 198 (99) | **0.0287*** | 24 (12) | 54 (27) | 122 (61) | **0.0244** | 31 (15.5) | 169 (84.5) | 0.1637 | 2 (28.6) | 22 (48.9) | 28 (36.4) | 148 (51) | **0.0401** | 135 (67.5) | 65 (32.5) | 0.6387 |
| | 3 | 152 | 6 (3.9) | 146 (96.1) | | 24 (15.8) | 30 (19.7) | 98 (64.5) | | 26 (17.1) | 126 (82.9) | | 5 (71.4) | 21 (46.7) | 24 (44.2) | 102 (35.2) | | 96 (63.2) | 56 (36.8) | |
| | *n* | *409* | *8 (2)* | *401 (98)* | | *50 (12.3)* | *93 (22.7)* | *266 (65)* | | *50 (12.2)* | *359 (87.8)* | | *7 (1.7)* | *45 (11)* | *67 (16.4)* | *290 (70.9)* | | *270 (66)* | *139 (34)* | |
| Age | ≥50 yr | 270 | 4 (1.5) | 266 (98.5) | | 30 (11.1) | 62 (23) | 178 (65.9) | | 44 (16.3) | 226 (83.7) | | 3 (42.9) | 27 (64.3) | 41 (64.1) | 199 (72.1) | | 182 (67.4) | 88 (32.6) | |
| | <50 yr | 119 | 4 (3.4) | 115 (96.6) | 0.2550~ | 17 (14.3) | 27 (22.7) | 75 (63) | 0.6708 | 24 (20.2) | 95 (79.8) | 0.3855~ | 4 (57.1) | 15 (35.7) | 23 (35.9) | 77 (27.9) | 0.1995 | 72 (61) | 46 (39) | 0.2464~ |
| | *n* | *389* | *8 (2.1)* | *381 (97.9)* | | *47 (12.1)* | *89 (22.9)* | *253 (65)* | | *68 (17.5)* | *321 (82.5)* | | *7 (1.8)* | *42 (10.8)* | *64 (16.5)* | *276 (70.9)* | | *254 (65.4)* | *134 (34.5)* | |
| Lymph node status | Negative | 95 | 3 (3.2) | 92 (96.8) | | 17 (17.9) | 21 (22.1) | 57 (60) | | 22 (23.2) | 73 (76.8) | | 3 (60) | 15 (51.7) | 20 (64) | 57 (49.1) | | 63 (67) | 31 (33) | |
| | Positive | 84 | 2 (2.4) | 82 (97.6) | >0.9999~ | 15 (17.9) | 19 (22.6) | 50 (59.5) | 0.9965 | 11 (13.1) | 73 (86.9) | 0.1215~ | 2 (40) | 14 (48.3) | 9 (36) | 59 (50.9) | 0.2787 | 56 (66.7) | 28 (33.3) | >0.9999~ |
| | *n* | *179* | *5 (2.8)* | *174 (97.2)* | | *32 (17.9)* | *40 (22.3)* | *107 (59.8)* | | *33 (18.4)* | *146 (81.6)* | | *5 (2.8)* | *29 (16.2)* | *29 (16.2)* | *116 (64.8)* | | *119 (66.9)* | *59 (33.1)* | |
| Tumor size | <2 cm | 134 | 1 (0.7) | 133 (99.3) | | 22 (16.4) | 28 (20.9) | 84 (62.7) | | 32 (23.9) | 102 (76.1) | | 1 (16.7) | 21 (58.3) | 30 (55.6) | 82 (45.3) | | 86 (64.7) | 47 (35.3) | |
| | 2-5 cm | 117 | 5 (4.3) | 112 (95.7) | 0.1002 | 15 (12.8) | 34 (29.1) | 68 (58.1) | 0.2427 | 23 (19.7) | 94 (80.3) | **0.0765*** | 5 (83.3) | 12 (33.3) | 22 (40.7) | 78 (43.1) | 0.1494 | 87 (74.4) | 30 (25.6) | 0.2533 |
| | >5 cm | 26 | 0 (0) | 26 (100) | | 3 (11.5) | 3 (11.5) | 20 (76.9) | | 2 (7.7) | 24 (92.3) | | 0 (0.0) | 3 (8.3) | 2 (3.7) | 21 (11.6) | | 18 (69.2) | 8 (30.8) | |
| | *n* | *277* | *6 (2.2)* | *271 (97.8)* | | *40 (14.4)* | *65 (23.5)* | *172 (62.1)* | | *57 (20.6)* | *220 (79.4)* | | *6 (2.2)* | *36 (13)* | *54 (19.5)* | *181 (65.3)* | | *191 (69.2)* | *85 (30.8)* | |
| Ki67 expression (20% threshold) | Low | 333 | 6 (1.8) | 327 (98.2) | | 45 (13.5) | 78 (23.4) | 210 (63.1) | | 55 (16.5) | 278 (83.5) | | 6 (85.7) | 40 (90.9) | 52 (78.8) | 235 (86.1) | | 221 (66.6) | 111 (33.4) | |
| | High | 57 | 2 (3.5) | 55 (96.5) | 0.3307~ | 4 (7.0) | 12 (21.1) | 41 (71.9) | 0.3092 | 16 (28.1) | 41 (71.9) | **0.0422~** | 1 (14.3) | 4 (9.1) | 14 (21.2) | 38 (13.9) | 0.3228 | 37 (64.9) | 20 (35.1) | 0.8796~ |
| | *n* | *390* | *8 (2.1)* | *382 (97.9)* | | *49 (12.6)* | *90 (23)* | *251 (64.4)* | | *71 (18.2)* | *319 (81.2)* | | *7 (1.8)* | *44 (11.3)* | *66 (16.9)* | *273 (70)* | | *258 (66.3)* | *131 (33.7)* | |
| ER status | Positive | 311 | 1 (0.3) | 310 (99.7) | | 34 (10.9) | 76 (24.4) | 201 (64.7) | | 62 (19.9) | 249 (80.1) | | 0 (0) | 34 (79.1) | 57 (85.1) | 220 (75.6) | | 210 (67.7) | 100 (32.3) | |
| | Negative | 97 | 7 (7.2) | 90 (92.8) | **0.0002~** | 14 (14.4) | 17 (17.5) | 66 (68.1) | 0.2936 | 10 (10.3) | 87 (89.7) | **0.0325~** | 7 (100) | 9 (20.9) | 10 (14.9) | 71 (24.4) | **<0.0001** | 59 (60.8) | 38 (39.2) | 0.2206~ |
| | *n* | *408* | *8 (2)* | *400 (98)* | | *48 (11.8)* | *93 (22.8)* | *267 (65.4)* | | *72 (17.6)* | *336 (82.4)* | | *7 (1.7)* | *43 (10.5)* | *67 (16.4)* | *291 (71.3)* | | *269 (66.1)* | *138 (33.9)* | |
| Subtypes | HR+/HER2- | 284 | 1 (0.4) | 283 (99.6) | | 32 (11.3) | 70 (24.6) | 182 (64.1) | | 57 (20.1) | 227 (79.9) | | 0 (0) | 32 (74.4) | 52 (77.6) | 200 (69) | | 191 (67.5) | 92 (32.5) | |
| | HER2+ | 50 | 0 (0) | 50 (100) | **<0.0001** | 5 (10) | 10 (20) | 35 (70) | 0.6395 | 5 (10) | 45 (90) | 0.1398 | 0 (0) | 5 (11.6) | 5 (7.5) | 40 (13.8) | **<0.0001** | 29 (58) | 21 (42) | 0.4256 |
| | TNBC | 73 | 7 (9.6) | 66 (90.4) | | 11 (15.1) | 13 (17.8) | 49 (67.1) | | 10 (13.7) | 63 (86.3) | | 7 (100) | 6 (14) | 10 (14.9) | 50 (17.2) | | 48 (65.8) | 25 (34.2) | |
| | *n* | *407* | *8 (2)* | *399 (98)* | | *48 (11.8)* | *93 (22.9)* | *266 (65.3)* | | *72 (17.7)* | *335 (82.3)* | | *7 (1.7)* | *43 (10.6)* | *67 (16.5)* | *290 (71.2)* | | *268 (66)* | *138 (34)* | |

### Figure S1

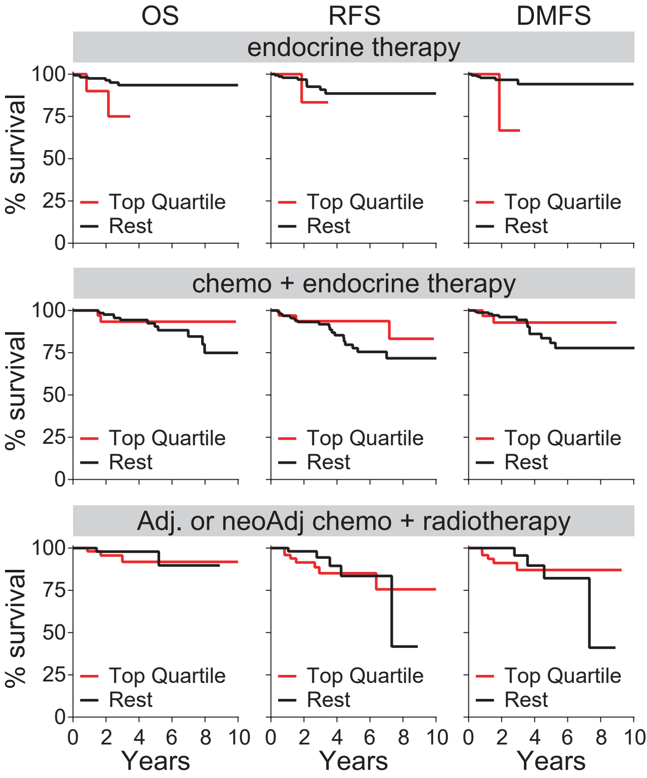
OS
RFS
DMFS
endocrine therapy
chemo + endocrine therapy
Adj. or neoAdj chemo + radiotherapy
% survival
100
75
50
25
0
Top Quartile
Rest
0
2
4
6
8
10
Years

### Figure S2

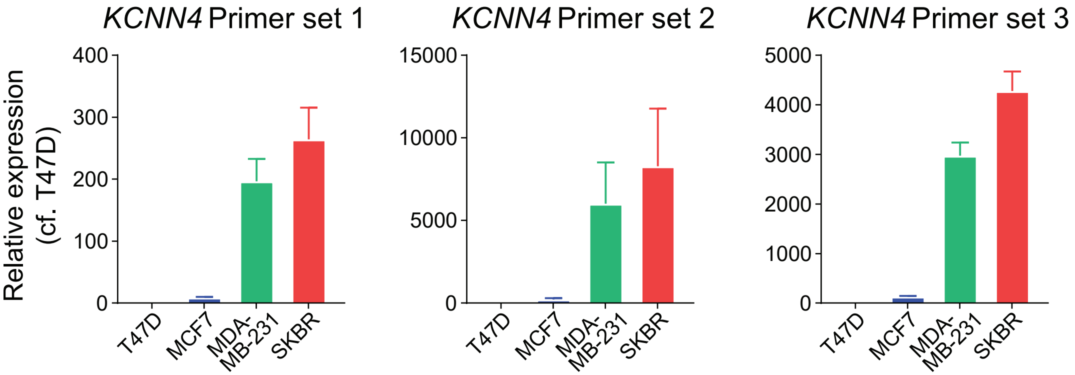

KCNN4 Primer set 1
KCNN4 Primer set 2
KCNN4 Primer set 3
Relative expression
(cf. T47D)
400
300
200
100
0
15000
10000
5000
0
5000
4000
3000
2000
1000
0
T47D
MCF7
MDA-
MB-231
SKBR

### Figure S3

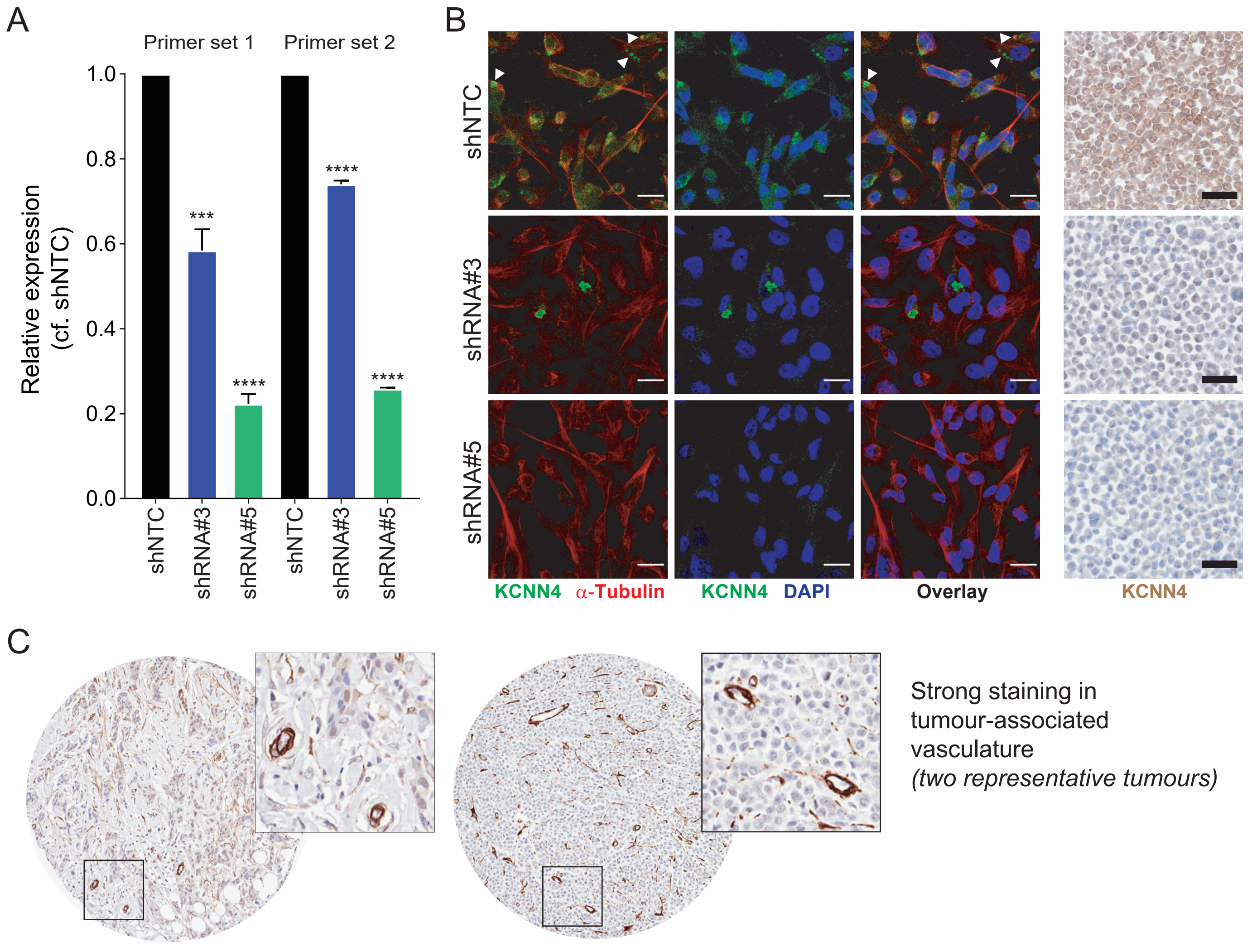

A
Primer set 1
Primer set 2
Relative expression (cf. shNTC)
1.0
0.8
0.6
0.4
0.2
0.0
***
****
****
****
shNTC
shRNA#3
shRNA#5
shNTC
shRNA#3
shRNA#5
B
shNTC
shRNA#3
shRNA#5
KCNN4 α-Tubulin
KCNN4 DAPI
Overlay
KCNN4
C
Strong staining in tumour-associated vasculature
(two representative tumours)

### Figure S4

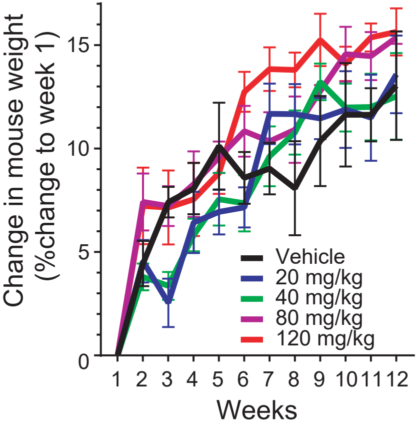

Change in mouse weight
(%change to week 1)
0
5
10
15
1
2
3
4
5
6
7
8
9
10
11
12
Weeks
Vehicle
20 mg/kg
40 mg/kg
80 mg/kg
120 mg/kg

### Figure S5

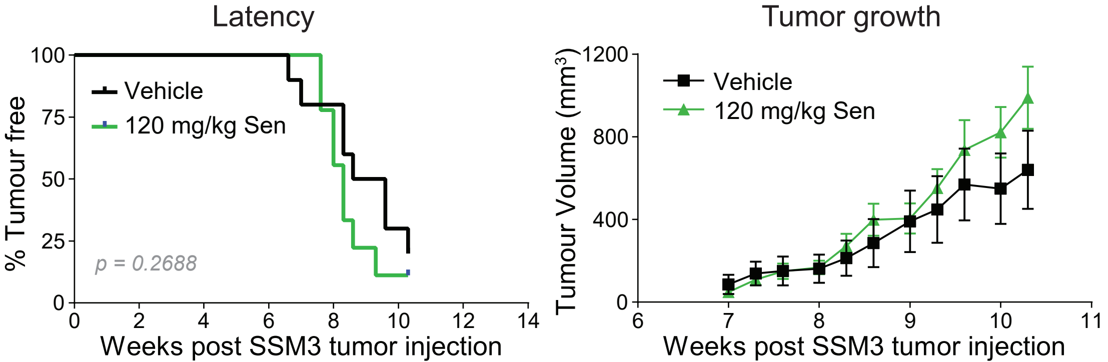

Latency
100
75
50
25
0
% Tumour free
Vehicle
120 mg/kg Sen
p = 0.2688
0
2
4
6
8
10
12
14
Weeks post SSM3 tumor injection
Tumor growth
1200
800
400
0
Tumour Volume (mm3)
Vehicle
120 mg/kg Sen
6
7
8
9
10
11
Weeks post SSM3 tumor injection

### Figure S6

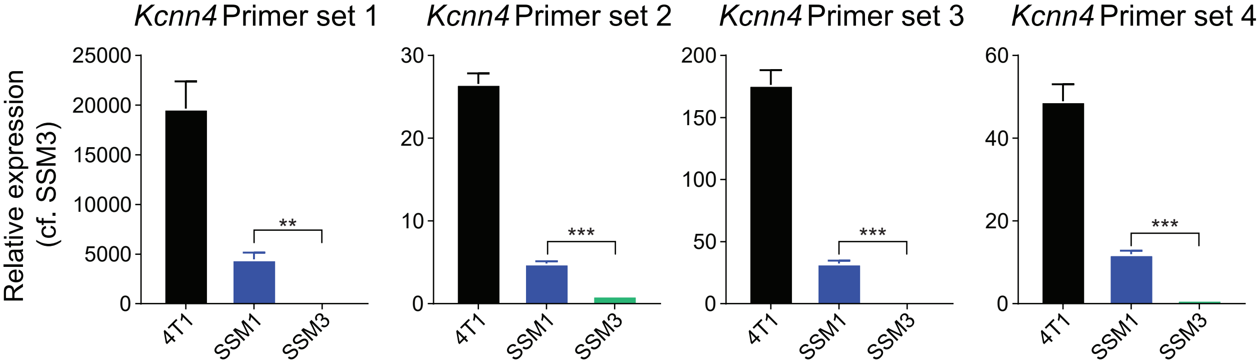

*Kcnn4* Primer set 1
*Kcnn4* Primer set 2
*Kcnn4* Primer set 3
*Kcnn4* Primer set 4
Relative expression (cf. SSM3)
25000
20000
15000
10000
5000
0
30
20
10
200
150
100
50
60
40
**
***
4T1
SSM1
SSM3

### Figure S7

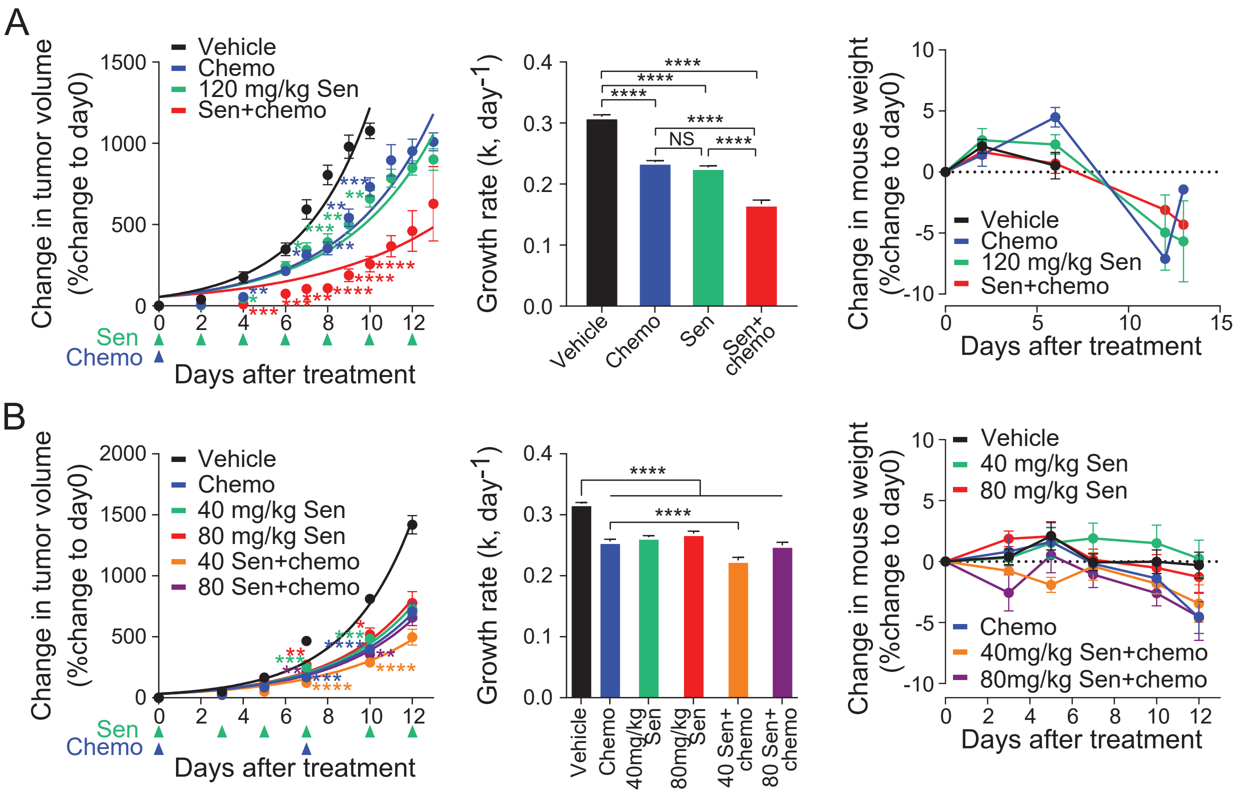

A
Vehicle
Chemo
120 mg/kg Sen
Sen+chemo
Change in tumor volume (%change to day0)
Days after treatment
Sen
Chemo
Growth rate (k, day$^{-1}$)
NS
Vehicle
Chemo
Sen
Sen+ chemo
Change in mouse weight (%change to day0)
Days after treatment
B
Vehicle
Chemo
40 mg/kg Sen
80 mg/kg Sen
40 Sen+chemo
80 Sen+chemo
Change in tumor volume (%change to day0)
Days after treatment
Growth rate (k, day$^{-1}$)
Vehicle
Chemo
40mg/kg Sen
80mg/kg Sen
40 Sen+ chemo
80 Sen+ chemo
Change in mouse weight (%change to day0)
40mg/kg Sen+chemo
80mg/kg Sen+chemo
Days after treatment

### Figure S8

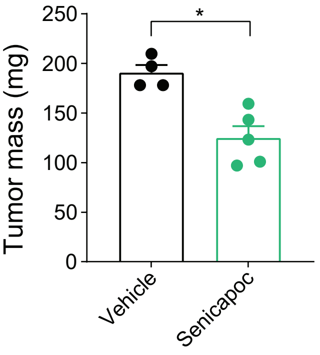

### Figure S9

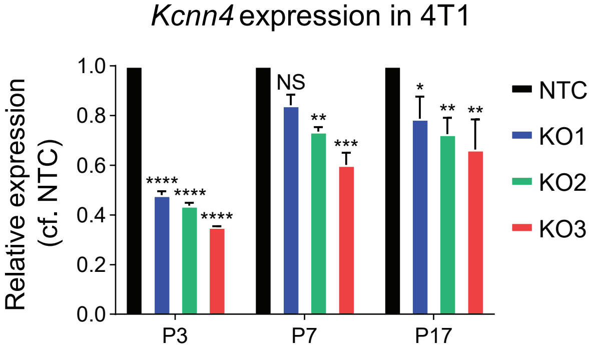
*Kcnn4* expression in 4T1
Relative expression (cf. NTC)
1.0
0.8
0.6
0.4
0.2
0.0
P3
P7
P17
NS
*
**
***
****
NTC
KO1
KO2
KO3

### Figure S10

#### Synthego - CRISPR Analysis on NTC clones

| SAMPLE | GUIDE TARGET | PAM SEQUENCE | INDEL % | MODEL FIT ($R^2$) | KNOCKOUT-SCORE |
|---|---|---|---|---|---|
| 4T1 NTC 2A8 | TGTGGGGCAAGATTGTCTGC | CTG | 0 | 1 | 0 |
| 4T1 NTC 1A4 | TGTGGGGCAAGATTGTCTGC | CTG | 0 | 1 | 0 |
| 4T1 NTC 1C2 | TGTGGGGCAAGATTGTCTGC | CTG | 0 | 1 | 0 |
| 4T1 NTC 1B12 | TGTGGGGCAAGATTGTCTGC | CTG | 0 | 1 | 0 |
| 4T1 NTC 1A8 | TGTGGGGCAAGATTGTCTGC | CTG | 0 | 1 | 0 |
| 4T1 NTC 1B5 | TGTGGGGCAAGATTGTCTGC | CTG | 0 | 1 | 0 |
| 4T1 NTC 1H5 | TGTGGGGCAAGATTGTCTGC | CTG | 0 | 1 | 0 |
| 4T1 NTC 2D11 | TGTGGGGCAAGATTGTCTGC | CTG | 0 | 1 | 0 |
| 4T1 NTC 1G10 | TGTGGGGCAAGATTGTCTGC | CTG | 0 | 1 | 0 |
| 4T1 NTC 2A4 | TGTGGGGCAAGATTGTCTGC | CTG | 0 | 1 | 0 |
| 4T1 NTC 2H2 | TGTGGGGCAAGATTGTCTGC | CTG | 0 | 1 | 0 |
| 4T1 NTC 1H9 | TGTGGGGCAAGATTGTCTGC | CTG | 0 | 1 | 0 |

#### Synthego - CRISPR Analysis on KO3 clones

| SAMPLE | GUIDE TARGET | PAM SEQUENCE | INDEL % | MODEL FIT ($R^2$) | KNOCKOUT-SCORE |
|---|---|---|---|---|---|
| 4T1 KC3 2A1 | TGTGGGGCAAGATTGTCTGC | CTG | 81 | 0.81 | 81 |
| 4T1 KC3 2B2 | TGTGGGGCAAGATTGTCTGC | CTG | 81 | 0.81 | 81 |
| 4T1 KC3 1E5 | TGTGGGGCAAGATTGTCTGC | CTG | 80 | 0.8 | 80 |
| 4T1 KC3 1H5 | TGTGGGGCAAGATTGTCTGC | CTG | 84 | 0.84 | 84 |
| 4T1 KC3 1C3 | TGTGGGGCAAGATTGTCTGC | CTG | 84 | 0.84 | 84 |
| 4T1 KC3 2C8 | TGTGGGGCAAGATTGTCTGC | CTG | 85 | 0.85 | 85 |
| 4T1 KC3 2B6 | TGTGGGGCAAGATTGTCTGC | CTG | 85 | 0.85 | 85 |
| 4T1 KC3 1E7 | TGTGGGGCAAGATTGTCTGC | CTG | 81 | 0.81 | 81 |
| 4T1 KC3 2A6 | TGTGGGGCAAGATTGTCTGC | CTG | 81 | 0.81 | 81 |
| 4T1 KC3 2C10 | TGTGGGGCAAGATTGTCTGC | CTG | 85 | 0.85 | 85 |
| 4T1 KC3 1A8 | TGTGGGGCAAGATTGTCTGC | CTG | 85 | 0.85 | 85 |
| 4T1 KC3 1E3 | TGTGGGGCAAGATTGTCTGC | CTG | 82 | 0.82 | 82 |

### Figure S11

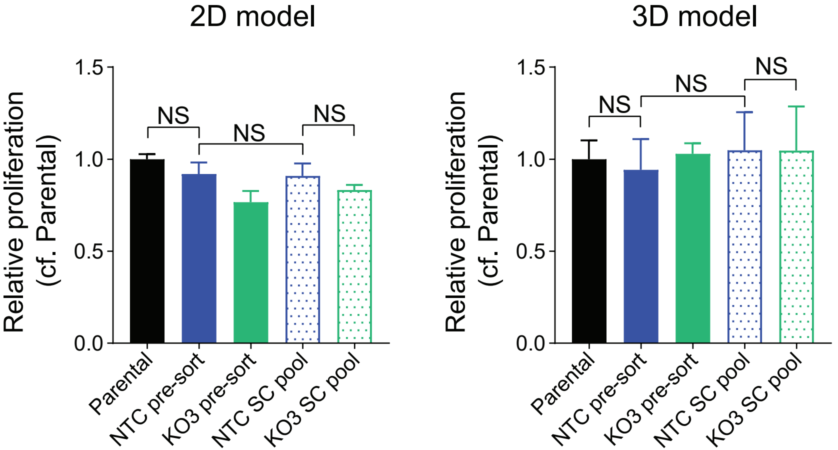

2D model
3D model
Relative proliferation
(cf. Parental)
1.5
1.0
0.5
0.0
NS
Parental
NTC pre-sort
KO3 pre-sort
NTC SC pool
KO3 SC pool

### Figure S12

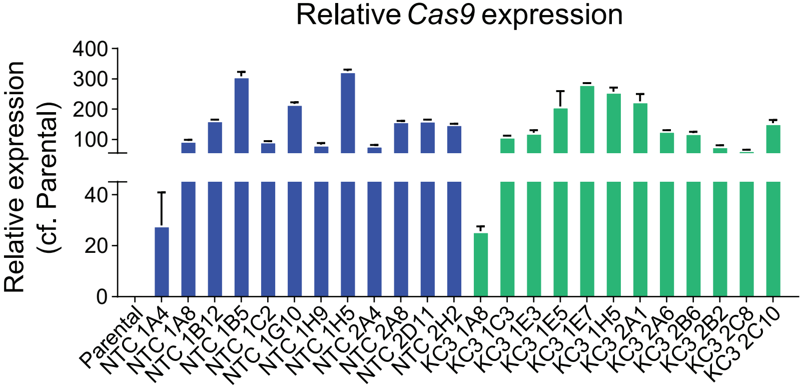
Relative *Cas9* expression
Relative exprestion
(cf. Parental)
400
300
200
100
40
20
0
Parental
NTC 1A4
NTC 1A8
NTC 1B12
NTC 1B5
NTC 1C2
NTC 1G10
NTC 1H9
NTC 1H5
NTC 2A4
NTC 2A8
NTC 2D11
NTC 2H2
KC3 1A8
KC3 1C3
KC3 1E3
KC3 1E5
KC3 1E7
KC3 1H5
KC3 2A1
KC3 2A6
KC3 2B6
KC3 2B2
KC3 2C8
KC3 2C10

### Figure S13

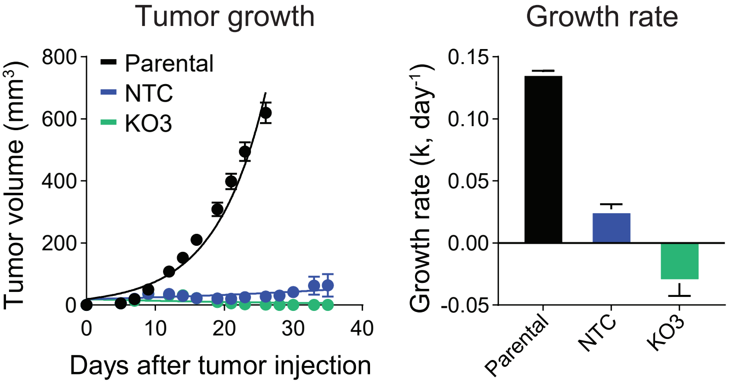
Tumor growth
Growth rate
800
600
400
200
0
Tumor volume (mm$^3$)
Parental
NTC
KO3
0
10
20
30
40
Days after tumor injection
0.15
0.10
0.05
0.00
-0.05
Growth rate (k, day$^{-1}$)
Parental
NTC
KO3
